# ppGpp Is Present in and Functions to Regulate Sleep in Drosophila

**DOI:** 10.1101/2021.09.16.460595

**Authors:** Way Young, Xiaohui Zhang, Huimin Daixi, Enxing Zhou, Ying Liu, Tao Wang, Wenxia Zhang, Xinxiang Zhang, Yi Rao

## Abstract

Discovery of molecules in living systems and demonstration of their functional roles are pivotal in furthering our understanding of the molecular basis of biology. ppGpp (guanosine-5’-diphosphate, 3’-diphosphate) has been detected in prokaryotes for more than five decades. Here we report that a genetic screen followed by chemical analysis revealed the presence of ppGpp in Drosophila. It can be detected in germ-free Drosophila and its level is controlled by an enzyme encoded by the *mesh1* gene in Drosophila. Loss of function mutations in *mesh1*, which encoded the ppGpp degrading enzyme led to longer sleep latency and less total sleep. These phenotypes could be rescued by wild type *mesh1*, but not by the enzymatically defective mutant Mesh1E66A, functionally implicating ppGpp. Ectopic expression of RelA, the *E. coli* synthetase for ppGpp, phenocopied *mesh1* knockout mutants, whereas overexpression of *mesh1* resulted in the opposite phenotypes, supporting that ppGpp is both necessary and sufficient in sleep regulation. *mesh1* is expressed in a specific population of neurons, and a chemoconnectomic screen followed by genetic intersection experiments implicate the pars intercerebralis (PI) as the site of ppGpp function. Our results have thus revealed that ppGpp is present in animals after long lag since its discovery in bacteria. Furthermore, we have demonstrated that ppGpp in a specific subset of neurons plays a physiological role in regulating sleep. We speculate that ppGpp may play function in mammals.

## INTRODUCTION

It was 52 years ago when guanosine-5’-diphosphate, 3’-diphosphate (guanosine tetraphosphate, ppGpp) and guanosine-5’-triphosphate, 3’-diphosphate (guanosine pentaphosphate, pppGpp) were implicated in gene regulation in *Escherichia coli* (*E. coli*) (Cashel and Gallant, 1969). Collectively known as (p)ppGpp, they are key players in bacterial stringent response to amino acid starvation (Dalebroux and Swanson, 2012; Field, 2018; Gourse *et al*., 2018; Hauryliuk *et al*., 2015; Liu, Nbittner and Wang, 2015; Magnusson, Farewell and Nyström, 2005; Potrykus *et al*., 2008; Wang, Sanders and Grossman, 2007). The level of ppGpp is regulated by the RelA/SpoT Homolog (RSH) family (Haseltine and Block, 1973; Hogg *et al*., 2004). In bacteria, RelA is one of the RSHs (Haseltine and Block, 1973), which contains both a ppGpp synthetase domain (SYNTH) and an inactive ppGpp hydrolase domain (HD) (Hogg *et al*., 2004). When amino acids are depleted, uncharged tRNAs accumulate (Fangman and Neidhardt, 1964) and activate the ribosome-associated RelA (Yang and Ishiguro, 2001), which increases the production of ppGpp (Cochran and Byrne, 1974). Another typical RSH in bacteria is the ppGpp hydrolase SpoT (Laffler and Gallant, 1974), containing a weak SYNTH (Leung and Yamazaki, 1997) and an active HD (Murry and Bremer, 1996). Upon iron limitation (Vinella *et al*., 2005), carbon starvation (Lesley and Shapiro, 2008), or glucose phosphate stress (Kessler *et al*., 2017), ppGpp was also increased (Potrykus and Cashel, 2008).

ppGpp has also been detected in plants (Boniecka *et al*., 2017; Ito *et al*., 2012; Kaisai *et al*., 2004; Sugliani *et al*., 2016; Tozawa *et al*., 2007; van der Biezen *et al*., 2000; Xiong *et al*., 2001). More than 30 members of the RSHs have been found in bacteria and plants (Atkinson *et al*., 2011; Field, 2018).

(p)ppGpp has not been detected in animals (Silverman and Atherly, 1979). Early report of ppGpp in the mouse turned out to be irreproducible (Irr, Kaulenas and Unsworth, 1974; Martini, Irr and Richter, 1977; Siverman and Atherly, 1977). Work in mammalian cell lines also failed to detect ppGpp before or after amino acid deprivation (Dabrowska *et al*., 2006a; Fan, Fisher and Edlin, 1973; Givens *et al*., 2004; Kim *et al*., 2009; Mamont *et al*., 1972; Rapaport and Bucher, 1976; Sato *et al*., 2015; Stanners and Thompson, 1974; Smulson, 1970; Takahashi *et al*., 2004; Thammana, Buerk and Gordon, 1976; Yamada *et al*., 2004). In Drosophila, Mesh1, a member of the RSH family containing only the hydrolase domain for ppGpp has been found (Sun *et al*., 2010), but ppGpp has not been detected.

Drosophila has served as a model for genetic studies of sleep for two decades (Hendricks *et al*., 2000, Shaw *et al*., 2000). Fly sleep is regulated by multiple genes functioning in several brain regions such as the fan-shape bodies (FSBs), the mushroom bodies, and the pars intercerebralis (PI) (e.g., Agosto *et al*., 2008; Andretic *et al*., 2005; Bushey *et al*., 2007; Cirelli *et al*., 2005; Crocker *et al*., 2010; Dai *et al*., 2019; Donlea *et al*., 2011, 2014; Guo *et al*., 2016; Hendericks *et al*., 2003; Koh *et al*., 2008; Kunst *et al*., 2014; Liu *et al*., 2014; Liu *et al*., 2016; Parisky *et al*., 2008; Qian *et al*., 2017; Shang *et al*., 2011; Sheeba *et al*., 2008; Sitaraman *et al*., 2015; Wu *et al*., 2010; Wu *et al*., 2014; Yuan *et al*., 2006; Yurgel *et al*., 2019).

We have carried out a genetic screen for genes involved in sleep regulation and discovered that ppGpp is present in flies and regulates sleep. *mesh1* is expressed in a specific population of neurons, and the effects of RelA and Mesh1 overexpression could be detected when they were expressed in neurons, but not in non-neuronal cells. Further dissection narrowed down the functional significance of *mesh1* expressing neurons in the PI. Thus, after 52 years of its discovery, ppGpp has finally been found in animals and shown to play an important physiological role in specific neurons.

## RESULTS

### Discovery of ppGpp in Drosophila

We screened through 1765 P-element insertion lines of Drosophila for mutations affecting sleep latency (Eddison *et al*., 2012) (Figure 1A). An insertion in the *mesh1* gene (*mesh1*-ins) (Figure 1C) was found to have significantly longer sleep latency (Figure 1A and 1D) and less total sleep duration (Figure 1 B).

**Figure 1.**
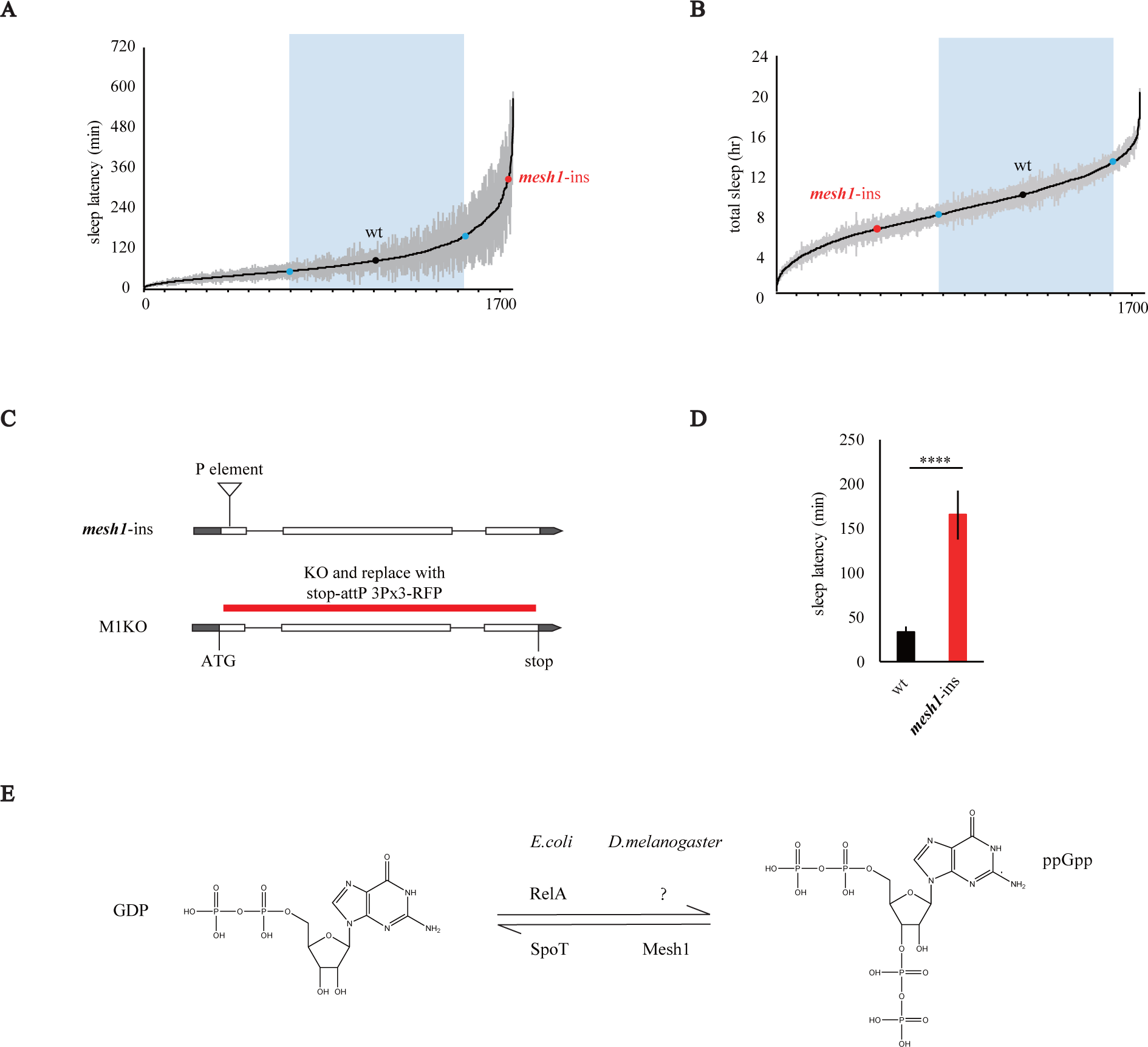
Mesh1 Gene and ppGpp Level in Drosophila. (A) Results of a screen of 1765 P-element insertion lines. The y axis shows sleep latency in minutes and the x axis shows P element insertion lines. Grey shadows showed the standard error of the mean (SEM) from multiple flies of each line, and blue dots and rectangles showed the range of 3-fold of standard deviation from the value of wt. *mesh1-ins* had longer sleep latency than the wt. (B) Results of a screen of 1765 P-element insertion lines. The y axis shows total sleep levels in hours and the x axis shows P element insertion lines. Grey shadows showed SEM from multiple flies of each line, and blue dots and rectangles showed the range of 3-fold of standard deviation from the value of wt. *mesh1-ins* had less total sleep than the wt. (C) Diagrams of *mesh1-*ins and M1KO. In *mesh1-*ins, the P-element was found to be inserted into the CDS of the 1^st^ exon. In M1KO, the entire CDS except the start codon was replaced with stop-2A-attP and 3Px3-RFP. Introns were labeled as straight lines, CDS as unfilled rectangles and untranslated regions (UTRs) as filled rectangles. (D) Statistical analysis of sleep latency. Compared to the wt, *mesh1-*ins exhibited significantly longer sleep latency (Student’s t-test, **** denotes p < 0.0001). (E) ppGpp metabolism. In *E. coli*, GDP is converted to ppGpp by its synthetase RelA, whereas ppGpp is converted to GDP by the hydrolase SpoT. In *D. melanogaster*, only the hydrolase Mesh1 has been discovered but the synthetase is unknown.

*mesh1* encodes an RSH family member in animals (Sun *et al*., 2010). Mesh1 protein is predicted to have hydrolase activity, converting ppGpp to GDP (Figure 1E). We expressed the Drosophila Mesh1 protein in *E. coli* and purified it. In an in vitro hydrolysis assay with ppGpp, Mesh1 indeed degraded ppGpp (Figure S1A).

ppGpp has not been detected in animals (Hauryliuk *et al*., 2015; Sun *et al*., 2010). To measure endogenous ppGpp in Drosophila, extracts from more than 2000 flies were made. ppGpp was detected by ultra-performance liquid chromatography and mass spectrometry (UPLC-MS) in wild type (wt) flies (Figure 2A, 2B, 2D, 2E).

**Figure 2.**
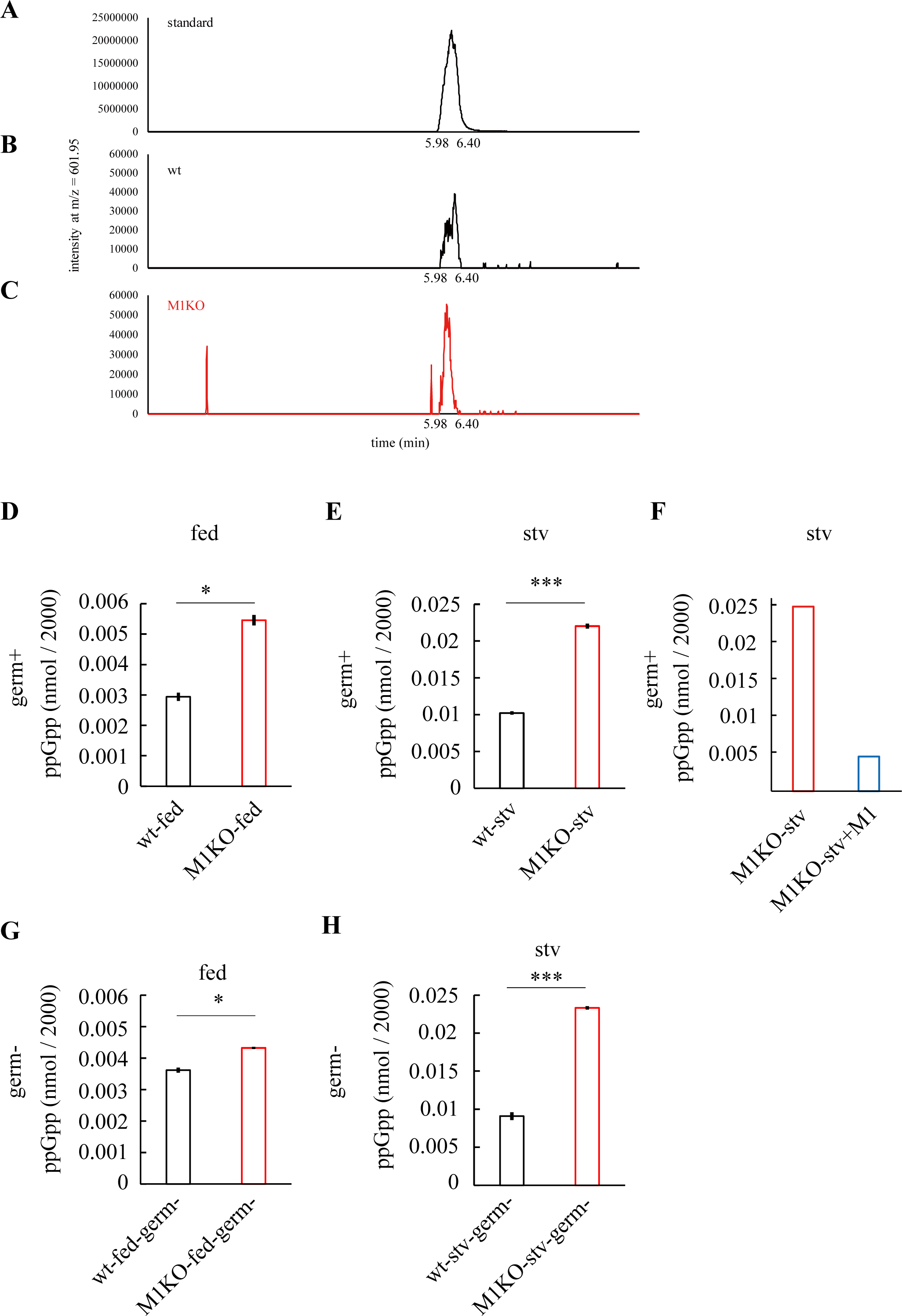
Identification and Quantitative Analysis of ppGpp in Drosophila. (A-C) UPLC-MS profiles of standard ppGpp (A), wt fly extracts (B), and M1KO fly extracts (C). Horizontal axis is time (minutes), and vertical axis is intensity of MS signal at m/z = 601.95, which corresponds to standard ppGpp (A). (D-E) Statistical analyses of ppGpp measurements under fed (D) or starved (E) conditions in *Drosophila*. For each genotype, approximately 2000 male flies were sacrificed for UPLC-MS measurements. In both fed and starved conditions, compared to that in wt, ppGpp level was significantly increased in M1KO (Student’s t-test, **** denoting p < 0.0001). (F) Mesh1 protein expressed in E.coli was purified and added into extracts from M1KO flies. ppGpp in the extracts was reduced by Mesh1. (G-H) Statistical analyses of ppGpp measurements under fed (D) or starved (E) conditions in germ-free Drosophila. Germ-free flies were generated and amplified (see method in details), and in each condition 2000 male flies were sacrificed for measurement by UPLC-MS.

We generated a knockout (KO) line of *mesh1* (M1KO) using CRISPR-Cas9 to replace most of its coding sequence following the start codon with 2A-attP (Figure 1C, S1B). More ppGpp was present in M1KO flies than the wt flies (Figure 2D and 2E), consistent with the fact that *mesh1* encoded only a hydrolase domain (Figure S1C). These results also supported that the ppGpp detected by us was regulated by the Drosophila *mesh1* gene.

The addition of Mesh1 protein to the extract from MIKO mutant flies also decreased the amount of ppGpp (Figure 2F).

Although ppGpp was detected and its level was regulated by the Drosophila *Mesh1* gene, which encoded an protein with ppGpp degrading activity, it remained possible that the detected ppGpp was synthesized from bacteria attached to flies. To test this possibility, we generated germ-free lines of wt and M1KO for analysis. We confirmed that the flies were indeed germ-free flies with both bacterial culture (Figure S1D) and 16S rDNA PCR (Figure S1E). ppGpp was detected in both fed and starved germ-free flies, and levels in M1KO were significantly higher in both conditions (Figure 2G, 2H).

Taken together, our results indicate that ppGpp exists in Drosophila.

### Sleep Phenotypes of M1KO Mutant Flies

M1KO null *mesh1* mutants allowed us to investigate *mesh1* function in sleep (Figure 3A). In the paradigm of 12 hours light and 12 hours darkness (LD), M1KO flies showed decreased nighttime sleep (Figure 3B), decreased total sleep (Figure 3D) and increased sleep latency at night (Figure 3E), but no change in day time sleep level (Figure 3C) or daytime sleep latency (Figure 3F). Sleep bout number and length either during the night or during the day were not significantly different between wt and M1KO flies (Figure S2A). Circadian rhythm was not significantly different between the wt and M1KO flies (Figure S2J, S2K, S2L and S2M). These results indicate that Mesh1 gene is mainly involved in regulating sleep latency.

**Figure 3.**
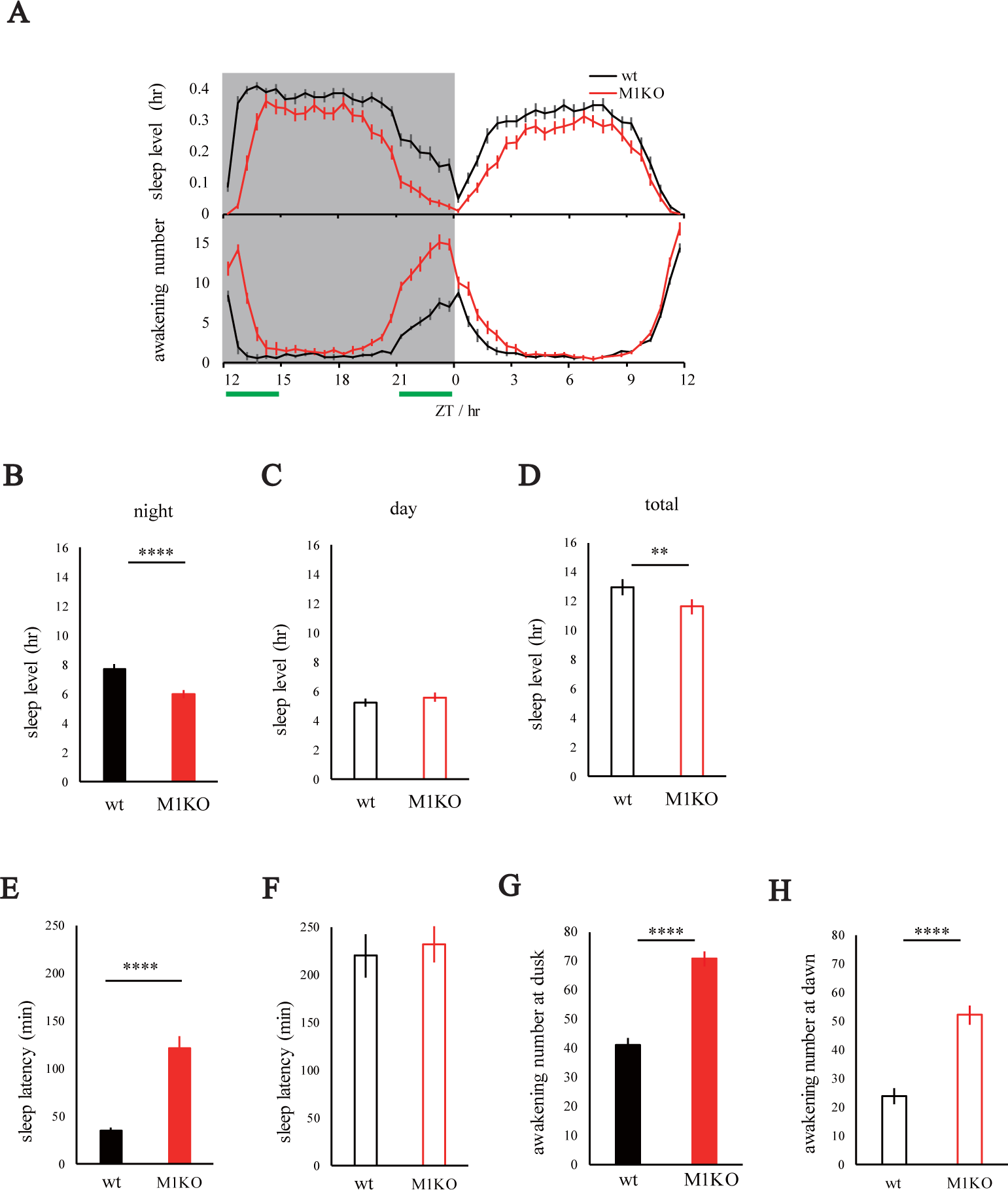
Sleep Phenotypes in M1KO Mutant Flies. (A) Profiles of sleep and awakening in M1KO Flies. Rest time for every 30 min was plotted (with the SEM as the error bars). Compared to the wt flies, M1KO flies began sleep later and wake up earlier. Dusk (ZT 0-3) and dawn (ZT 9-12) of night time sleep were denoted with green bars. (B-D) Statistical analyses of sleep level during night time (B), day time (C) or in total (D). Compared to the wt, M1KO flies slept significantly less at night (Student’s t-test, **, **** denoting p < 0.01, and p < 0.0001 respectively). (E-F) Statistical analyses of sleep latency during night time (E), or day time (F). Compared to the wt, M1KO flies exhibited a significant increase in night sleep latency (Student’s t-test, **** denoting p < 0.0001). (G-H) Statistical analyses of summed awakening numbers at dusk (G, ZT0-3) and dawn (H, ZT9-12) shown in (A). Awakening numbers of both dusk and dawn were significantly increased in M1KO flies (Student’s t-test, **** denoting p < 0.0001).

We assessed awakening according to a procedure reported recently (Tabuchi *et al*., 2018). Awakening number at the beginning of nighttime sleep (dusk) (Figure 3G) and awakening number near the end of nighttime sleep (dawn) (Figure 3H) were significantly increased in M1KO mutants.

Under the constant darkness (DD) condition, the same phenotypes in sleep level (Figure S2B, S2D, S2E), latency (Figure S2F, S2G) and awakening (Figure S2H and S2I) were observed. Sleep level in presumptive day is decreased in W1KO flies under DD (Figure S2D), but not under LD condition (Figure 3C).

### Requirement of the Enzymatic Activity in *mesh1* for Its Rescue of Sleep Phenotypes in M1KO Mutants

We generated another KO line of *mesh1* (M1KOGal4) by replacing its CDS after the start codon with an in-frame fusion of the 2A peptide and the yeast transcription factor Gal4 (Figure 4A and S1B). To examine whether *mesh1* gene functions in sleep through its regulation of ppGpp level, we compared the activities of the wt Mesh1 protein with that of the Mesh1 E66A mutant protein, both in vitro and in vivo.

**Figure 4.**
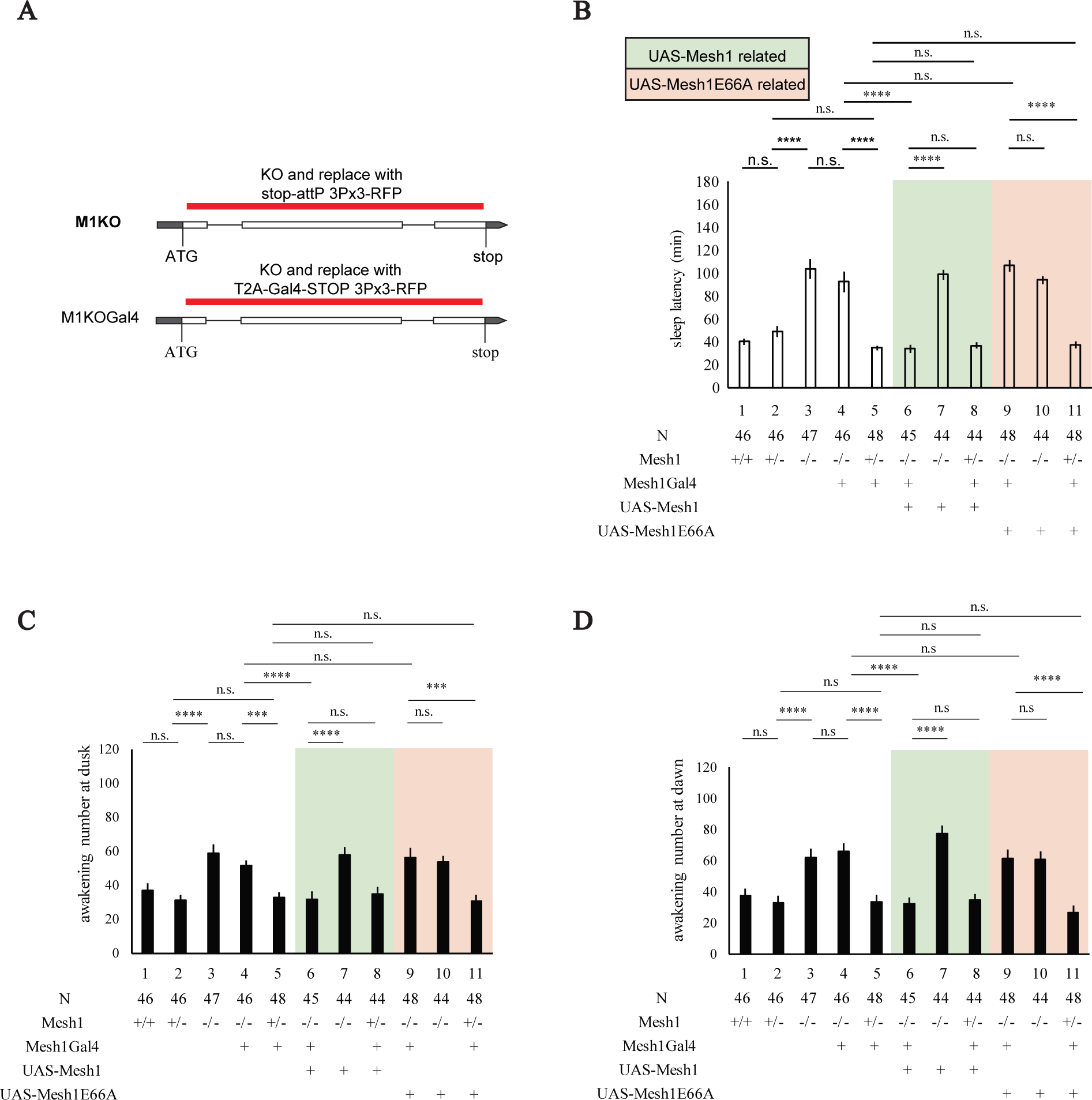
Significance of the Hydrolysis Activity of Mesh1 in Drosophila Sleep. (A) Diagrams of constructs used to generate M1KO and M1KOGal4 flies. *mesh1* CDS except the start codon was deleted in both M1KO and M1KOGal4, and the only difference is that a T2A-fused Gal4 (with a stop codon) were placed in M1KOGal4. (B) Statistical analyses of sleep latency at night. Columns 1-3: wt, M1KOGal4/+ and M1KOGal4/M1KOGal4 flies, with the sleep latency significantly increased in M1KOGal4/M1KOGal4 (Column 3). Columns 4-5: M1KOGal4/M1KO and M1KOGal4/+. Columns 6-8: M1KOGal4/M1KO > UAS-Mesh1/+, M1KO/M1KO, UAS-Mesh1/+, M1KOGal4/+>UAS-Mesh1/+. M1KOGal4/M1KO>UAS-Mesh1 (Column 6) could fully rescue the sleep latency phenotype (Columns 7-8). Columns 9-11: M1KOGal4/M1KO>UAS-Mesh1E66A/+, M1KO/M1KO, UAS-Mesh1E66A/+, M1KOGal4/+>UAS-Mesh1E66A/+. The hydrolysis-deficient line UAS-Mesh1E66A/+ driven by M1KOGal4/M1KO (Column 9) failed to rescue sleep latency (Columns 10-11). Comparison was based on two-way ANOVA and Bonferroni post-tests with **** denotes p < 0.0001. (C) Statistical analyses of awakening number at dusk. Columns 1-3: awakening number significantly increased in M1KO (Column 3). Columns 4-5: M1KOGal4/M1KO (Column 4) had more awakening number than the wt. Columns 6-8: M1KOGal4/M1KO > UAS-Mesh1 (Column 6) could rescue the awakening number deficiency (Columns 7-8). Columns 9-11: the hydrolysis-deficient UAS-Mesh1E66A/+ driven by M1KOGal4/M1KO (Column 9) failed to rescue the awakening number deficiency (Columns 10-11). (D) Statistical analyses of awakening number at dawn.

Mesh1E66A expressed in bacteria could not hydrolyze ppGpp in vitro (Figure S1A). Similar to that in M1KO mutants, the level of ppGpp in flies was also increased in M1KOGal4 (Figure S1G). ppGpp level was reduced to that of the wt flies when the wt *mesh1* gene was expressed in flies under the control of the M1KOGal4 driver (Figure S1G). By contrast, expression of *mesh1E66A* could not reduce the level of ppGpp in M1KOGal4 flies (Figure S1G). These results indicate that wt Mesh1 protein could, but Mesh1 E66A mutant protein could not, hydrolyze ppGpp either in vitro or in vivo.

M1KOGal4/M1KO flies were phenotypically similar to M1KO/M1KO in having longer sleep latencies (Figure 4B) and more awakening numbers (Figure 4D). Expression of UAS-*mesh1* driven by M1KOGal4 in the background of M1KOGal4/M1KO rescued all these phenotypes (Figure 4B to 4D). By contrast, UAS-*mesh1E66A* could not rescue the sleep phenotypes in M1KOGal4/M1KO flies (Figure 4B to 4D).

Taken together, the in vitro results from bacterially expressed Mesh1 and Mesh1E66A proteins and the in vivo results from genetic rescue experiments strongly support that ppGpp regulates sleep.

### Pattern of *mesh1* Expression in Drosophila

To examine the expression pattern of the *mesh1* gene, we crossed M1KOGal4 with each of the following four UAS lines: UAS-mCD8-GFP for neuronal overviews (Lee and Luo, 1999), UAS-redStinger for nuclei (Barolo, Castro and Posakony, 2004), UAS-denmark for dendrites (Nikolai *et al*., 2010) and UAS-syt.eGFP for axon terminals (Zhang, Rodesch and Broadie, 2002).

*mesh1* was found in neurons in the brain and the ventral nerve cord (Figure 5A and 5E). In the brain, *mesh1* expressing neurons were detected in the PI and the suboesophageal ganglia (SOG) (Figure 5A and 5B). The *mesh1* expressing neurons in the PI and their axonal terminals in the SOG (Figure 5A, 5C and 5D) were reminiscent of the insulin-producing cells (IPC) in the PI which were shown previously to regulate sleep (Crocker *et al*., 2010). Immunostaining with an antibody for the *Drosophila* insulin like peptide 2 (Dilp2) confirmed expression of *mesh1* in IPC neurons positive in the PI (Figure S4C and S4E) (Li and Gong, 2015).

**Figure 5.**
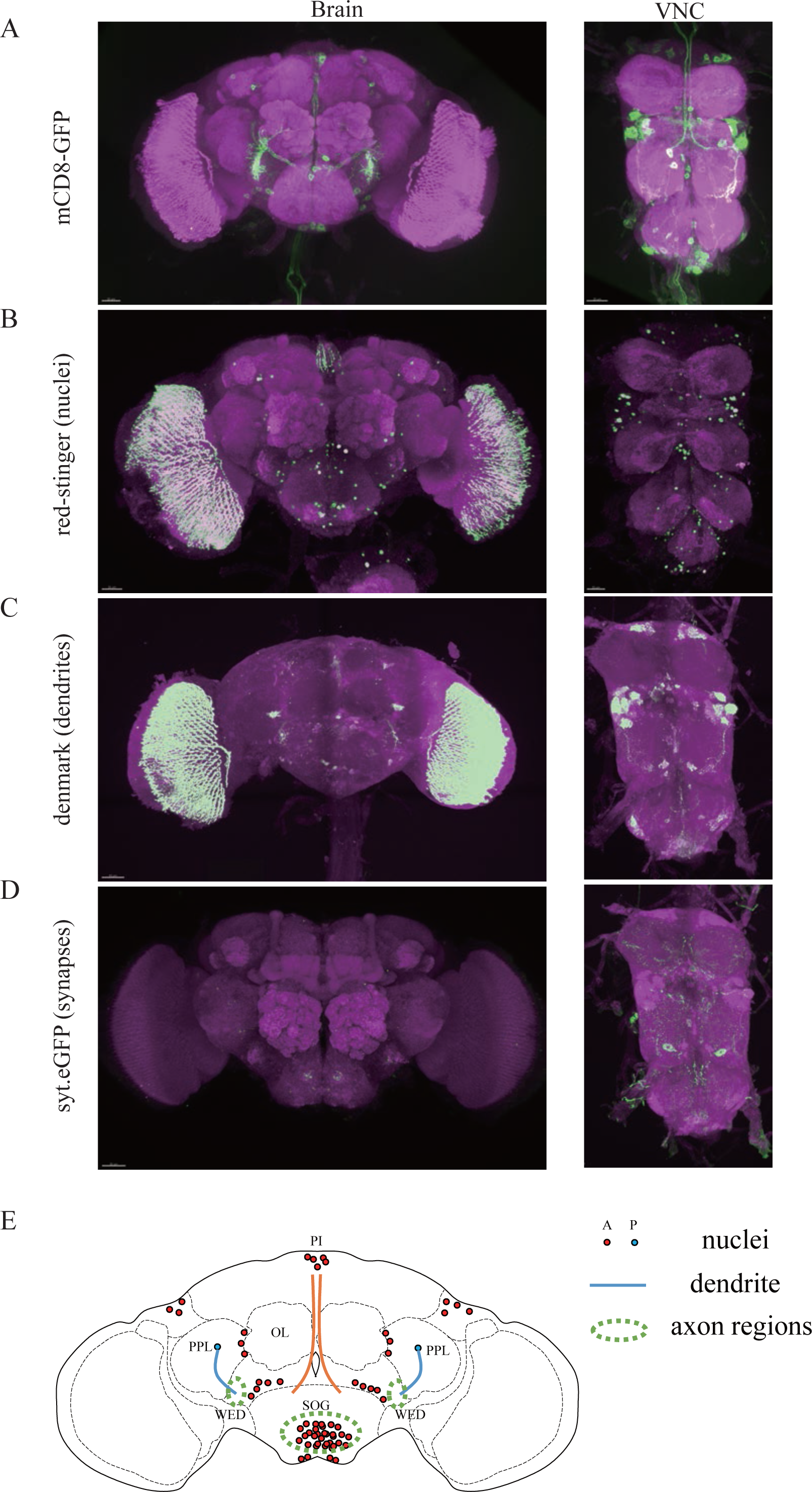
The Expression Pattern of M1KOGal4. (A) Neurons expressing mCD8-GFP driven by M1KOGal4 in the brain (left) and the VNC (right). Magenta showed immunostaining by the nc82 monoclonal antibody specific for neurons. (B) Nuclei expressing red-stinger driven by M1KOGal4 in the brain (left) and the VNC (right). Redstinger is presented in green. (C) Dendrites revealed by M1KOGal4>UAS-denmark in the brain (left) and the VNC (right). The dendritic marker denmark is presented in green. (D) Synaptic terminals revealed by M1KOGal4>UAS-Syt.eGFP in the brain (left) and the VNC (right). Syt.eGFP is presented in green. (E) A diagrammatic summary of neurons revealed by M1KOGal4. A: anterior sections. P: posterior sections.

### Evidence for ppGpp Functioning in Neurons

Because ppGpp regulate bacterial transcription through molecules such as the RNA polymerase ((Cashel and Gallant, 1969; Dalebroux and Swanson, 2012; Gourse *et al*., 2018; Hauryliuk *et al*., 2015; Magnusson, Farewell and Nyström, 2005; Potrykus *et al*., 2008; Wang, Sanders and Grossman, 2007), it is possible that ppGpp functions in many, even possibly all cells in animals. To test whether ppGpp functions in specific cells, we increased and decreased ppGpp levels in different populations of cells, by using ppGpp synthesizing and hydrolyzing enzymes.

The *E. coli RelA* gene encodes a synthetase for ppGpp (Laffler and Gallant, 1974). We confirmed that the RelA from *E. coli* used by us indeed increased ppGpp in vitro (Figure S1A). The biochemical activity of RelA protein is opposite to that of the Mesh1 protein, the effect of RelA overexpression in flies should be similar to *mesh1* knockout mutation. In flies with UAS-RelA driven either by tub-Gal4 for expression in all cells (O’Donnell *et al*., 1994) or by elav-Gal4 for expression in neurons (Robinow and White, 1991), sleep latency (Figure 6A) and awakening number were increased (Figure 6B). By contrast, UAS-RelA driven by repo-Gal4 for expression in glial cells (Halter *et al*., 1995) did not affect sleep (Figure 6A and 6B). These results indicate that ppGpp regulation of sleep does not involve glial cells, but specifically neurons.

**Figure 6.**
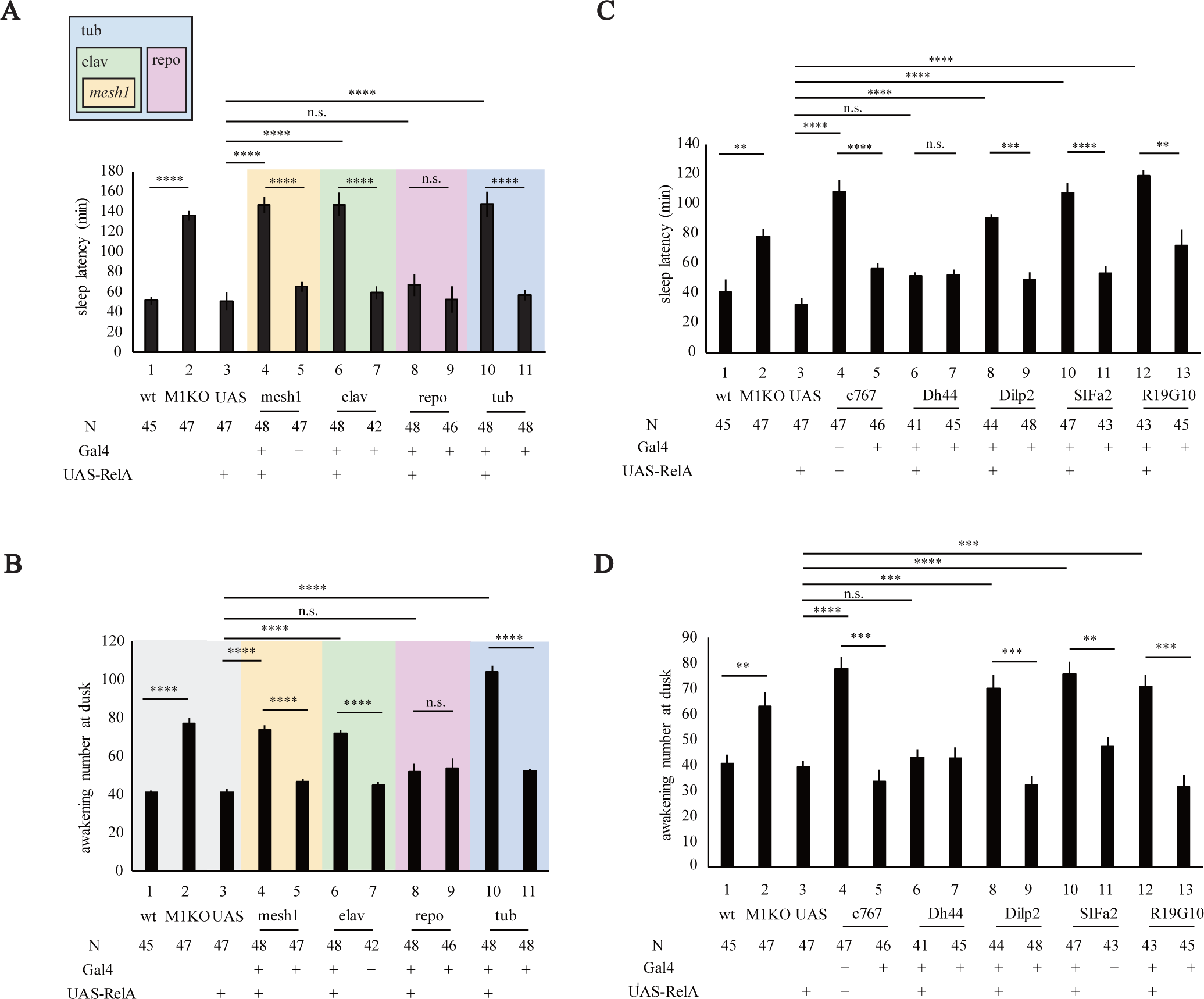
Effects of RelA Expression on Drosophila Sleep. (A) Statistical analyses of sleep latency when RelA was expressed in all cells with (tub-Gal4), neurons (elav-Gal4), glia (repo-Gal4) or mesh1 positive cells (mesh1Gal4). (1) wt , (2) M1KO, (3) UAS-RelA/+, (4) M1KOGal4/+>UAS-RelA/+, (5) M1KOGal4/+, (6) elav-Gal4/+> UAS-RelA/+, (7) elav-Gal4/+, (8) repo-Gal4/+>UAS-RelA/+, (9) repo-Gal4/+, (10) tub-Gal4/+>UAS-RelA/+, (11) tub-Gal4/+. Comparisons were based on two-way ANOVA and Bonferroni post-tests with **** denoting p < 0.0001. (B) Statistical analyses of awakening number at dusk when RelA was expressed in all cells (10), neuron (6), glia (8) or *mesh1* positive cells (4). (C) Statistical analyses of sleep latency with RelA expression in different subsets of PI neurons. (1) wt, (2) M1KO, (3) UAS-RelA/+, (4) c767/+>UAS-RelA/+, (5) c767/+, (6) Dh44-Gal4/+>UAS-RelA/+, (7) Dh44-Gal4/+, (8) Dilp2-Gal4/+> UAS-RelA/+, (9) Dilp2-Gal4/+, (10) SIFa2-Gal4/+>UAS-RelA/+, (11) SIFa2-Gal4/+, (12) R19610-Gal4/+>UAS-RelA/+, (13) UAS-RelA/+. Comparisons were based on two-way ANOVA and Bonferroni post-tests, with **, ***, **** denoting p < 0.01, p < 0.001, p < 0.0001 respectively. (D) Statistical analyses of awakening number with RelA expression in different subsets of PI neurons. Comparisons were based on two-way ANOVA and Bonferroni post-tests with **, ***, **** denoting p < 0.01, p < 0.001, p < 0.0001 respectively.

Furthermore, when M1KOGal4 was used to drive UAS-RelA expression, both sleep latency (Figure 6A) and awakening numbers (Figure 6B) were significantly increased, indicating that ppGpp level in *mesh1* positive neurons is sufficient to regulate sleep.

To decrease ppGpp level in fly cells, we used UAS-*mesh1*. UAS-*mesh1* overexpression driven by tub-Gal4 in all cells or by elav-Gal4 in neurons significantly decreased sleep latency (Figure 7A) and awakening numbers (Figure 7B). By contrast, UAS-*mesh1* overexpression driven by repo-Gal4 in glial cells did not affect sleep latency or awakening numbers (Figure 7A and 7B). When UAS-*mesh1* was driven by Mesh1 Gal4, sleep latency and awakening number were decreased (Figure 7A and 7B).

**Figure 7.**
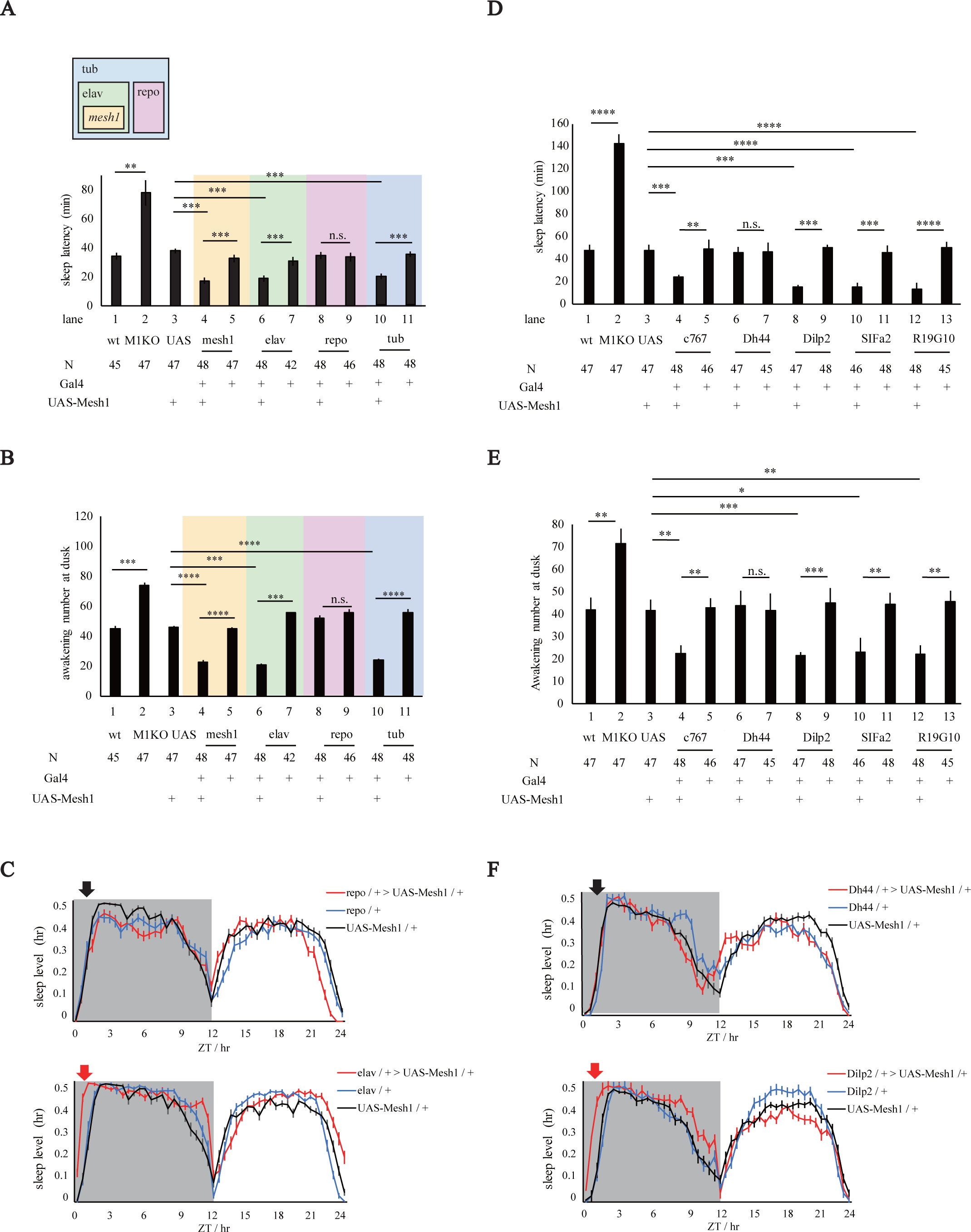
Effects of Mesh1 Overexpression on Drosophila Sleep. (A) Statistical analyses of sleep latency when Mesh1 was overexpressed in all cells, neurons, glia or *mesh1* positive cells. (1) wt, (2) M1KO, (3) UAS-Mesh1/+, (4) M1KOGal4/+ > UAS- Mesh1/+, (5) M1KOGal4/+, (6) elav-Gal4/+ > UAS- Mesh1/+, (7) elav-Gal4/+, (8) repo-Gal4/+>UAS-Mesh1/+, (9) repo-Gal4/+, (10) tub-Gal4/+, UAS-Mesh1/+, (11) tub-Gal4/+. Comparisons were based on two-way ANOVA and Bonferroni post-tests with **, *** denoting p < 0.01 and p < 0.001 respectively. (B) Statistical analyses of awakening number when Mesh1 was overexpressed in all cells, neurons, glia or *mesh1* positive cells. Comparisons were based on two-way ANOVA and Bonferroni post-tests with *** and **** denoting p < 0.001 and p < 0.0001 respectively. (C) Representative sleep profiles for lines in (A) and (B). Red arrow denotes sleep latency at dusk. UAS-Mesh1/+ in the top and bottom panels was the shared control, with the top panel showing that repo-Gal4>UAS Mesh1 did not affect sleep latency whereas the bottom panel showing that elav-Gal4>UAS Mesh1 shortened sleep latency. (D) Statistical analyses of sleep latency with Mesh1 overexpression in different PI subsets. (1) wt, (2) M1KO, (3) UAS-Mesh1/+, (4) c767-Gal4/+ and UAS-Mesh1/+, (5) c767-Gal4/+, (6) Dh44-Gal4/+ and UAS-Mesh1/+, (7) Dh44-Gal4/+, (8) Dilp2-Gal4/+ and UAS-Mesh1/+, (9) Dilp2-Gal4/+, (10) SIFa2-Gal4/+ and UAS-Mesh1/+, (11) SIFa2-Gal4/+, (12) R19G10-Gal4/+ and UAS-Mesh1/+, (13) R19G10-Gal4/+. Comparisons were based on two-way ANOVA and Bonferroni post-tests with **, ***, **** denoting p < 0.01, p < 0.001 and p < 0.0001 respectively. (E) Statistical analyses of awakening number with Mesh1 overexpression in different PI subsets. Comparisons were based on two-way ANOVA and Bonferroni post-tests with *, **, *** denoting p < 0.05, p < 0.01 and p < 0.001 respectively. (F) Representative sleep profiles for lines in (D) and (E). Red arrow denotes sleep latency at dusk. Note that the UAS-Mesh1/+ profile in the top and bottom panels was the shared control, with the top panel showing that Dh44-Gal4>UAS Mesh1 did not affect sleep latency whereas the bottom panel showing that Dilp2-Gal4>UAS Mesh1 shortened sleep latency.

Taken together, data from increasing or decreasing the ppGpp level by RelA and Mesh1 indicate that ppGpp functions in neurons but not in glia to regulate sleep.

### Dissection of PI Neurons Involved in ppGpp Regulation of Sleep

Our results have shown that neurons, specifically, *mesh1* expressing neurons, are required for ppGpp regulation of sleep (Figure 4B, 4C, 4D, 6A, 6B, 7A and 7B). We have generated a Gal4 library for the chemoconnectome (CCT), which include all the known neurotransmitters, modulators, neuropeptides and their receptors (Deng *et al*., 2019). We carried out a CCT screen by crossing each Gal4 line with UAS-RelA. RelA expression driven by Gal4 lines for Trh, Capa-R, CCHa2-R, LkR, OA2 and CG13229 affected sleep (Figure S3A). It was noted that all of these lines drove expression in the PI (Figure S3 B to G). This suggests functional significance of *mesh1* expression in the PI (Figure 5).

To further dissect the PI neurons for functional involvement in ppGpp regulation of sleep, we tested five Gal4 lines known to drive expression in PI neurons: a line for PI (c767, Cavanaugh *et al*., 2014; Donlea *et al*., 2009), three lines for neuropeptides Dh44 (Cannel *et al*., 2016; Chen and Dahanukar, 2018), Dilp2 (Crocker *et al*., 2010; Semaniuk *et al*., 2018; Yurgel *et al*., 2019) and SIFa (Martelli *et al*., 2017; Park *et al*., 2014), and one line for receptor R19G10 (an RYamide receptor) (Collins *et al*., 2011; Ohno *et al*., 2017). We crossed each of these PI drivers to a line of knockin flies with in-frame fusion of the flippase gene to the C terminus of the *mesh1* gene (M1KIflp) (Figure S4A), which was recombined with UAS-FRT-stop-FRT-mCD8-GFP as a reporter. Expression of *mesh1* gene in these specific PI neurons were confirmed (Figure S4B, S4C and S4D).

Each of these PI drivers was crossed with UAS-RelA to increase ppGpp level in specific neurons. Sleep latency (Figure 6C) and awakening (Figure 6D) were increased by RelA expression (Figure 6C and 6D) in neurons expressing Dilp2, SIFa, R19G10 or c767. However, RelA expression in Dh44 neurons could affect sleep (Figure 6C and 6D) Each of these PI drivers was crossed with UAS-Mesh1 to decrease ppGpp level in specific neurons. Overexpression of Mesh1 in Dilp2 neurons decreased sleep latency and awakening (Figure 7D, 7E and 7F). Sleep latency and awakening were decreased by Mesh1 expression in neurons positive for Dilp2, SIFa, R19G10 or c767 (Figure 7D and 7E). However, Mesh1 expression in Dh44 neurons could affect sleep (Figure 7D and 7E).

Taken together, experiments with RelA and Mesh1 expression have provided consistent results indicating that specific subsets of neurons in the PI are involved in ppGpp regulation of sleep.

### Role of ppGpp in Starvation Induced Sleep Loss

We investigated whether ppGpp played a role in starvation induced sleep loss (SISL). Similar to humans (MacFadyen, Oswald and Lewis, 1973), flies also show SISL (Keene *et al*., 2010). Baseline sleep in fed flies was recorded before flies were starved for 24 hours (Yurgel *et al*., 2019). Sleep during starvation was compared to that before starvation (Figure 8A).

**Figure 8.**
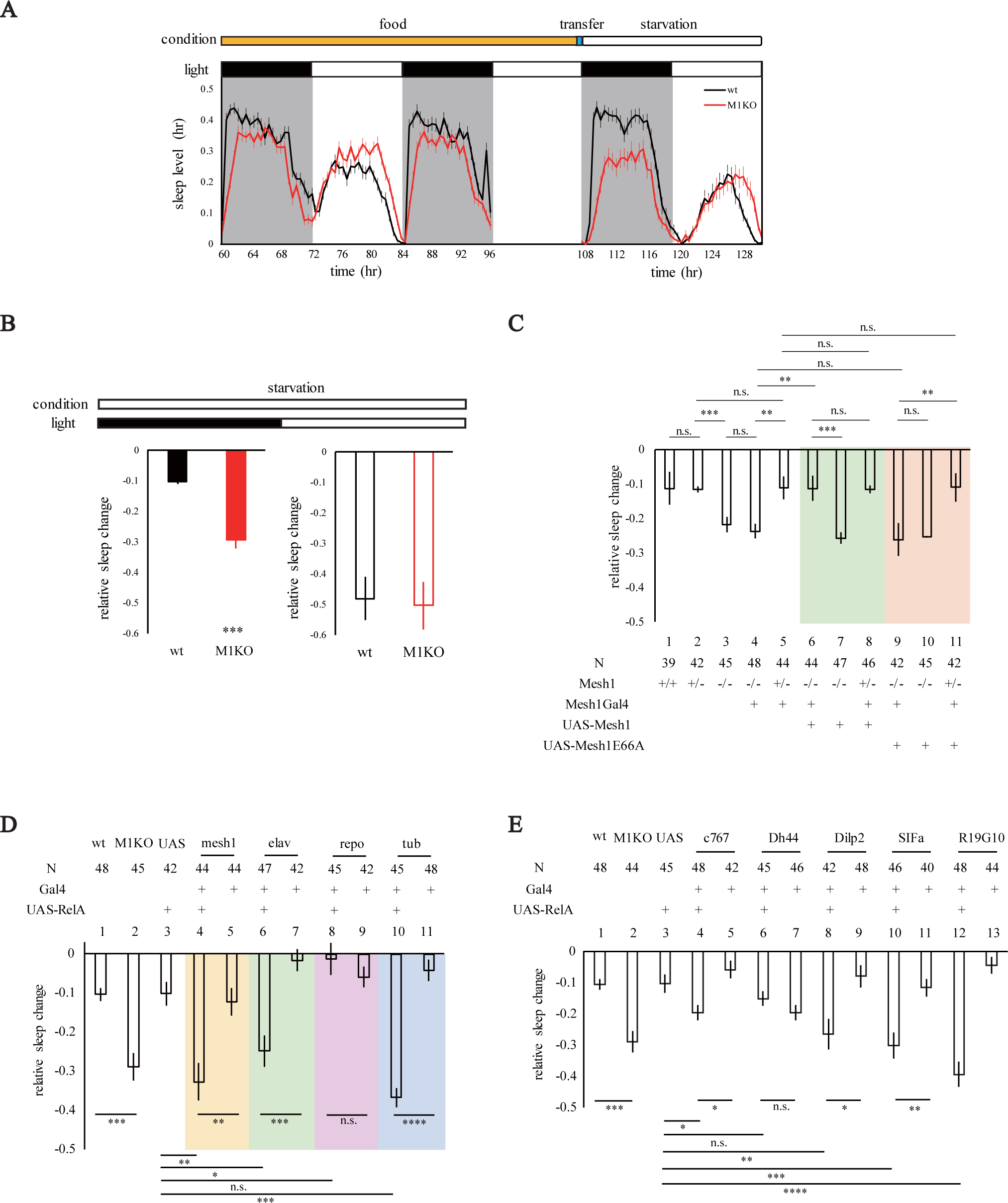
Effects of Mesh1 Knockout Mutations or RelA Expression on SISL. (A) SISL in M1KO. Baseline sleep was recorded on food for 3 days (recording at day 2 and day 3), before flies were transferred to 1% agar at the end of the 3rd day, followed by sleep recording while starvation of 24hr. (B) Statistical analysis of SISL in (A). Sleep was lost ∼10% at night and ∼50% at day, compared to the baseline in wt. This was significantly exacerbated in M1KO at night (Student’s t-test with *** denoting p < 0.001). (C) Statistical analyses of SISL with Mesh1 and Mesh1E66A rescue experiments. (1) wt, (2) M1KO/+, (3) M1KO/M1KO, (4) M1KOGal4/M1KO, (5) M1KOGal4/+, (6) M1KOGal4/M1KO and UAS-Mesh1/+, (7) M1KO/M1KO and UAS-Mesh1/+, (8) M1KOGal4/+ and UAS-Mesh1/+, (9) M1KOGal4/M1KO and UAS-Mesh1E66A/+, (10) M1KO/M1KO and UAS-Mesh1E66A/+, (11) M1KOGal4/+ and UAS-Mesh1E66A/+. Comparison was based on two-way ANOVA and Bonferroni post-tests with **, *** denoting p < 0.01, p < 0.001 respectively. (D) Statistical analyses of SISL with RelA expression in all cells, neurons, glia or *mesh1* positive cells. (1) wt, (2) M1KO, (3) UAS-RelA/+, (4) M1KOGal4/+> UAS-RelA/+, (5) M1KOGal4/+, (6) elav-Gal4/+>UAS-RelA/+, (7) elav-Gal4/+, (8) repo-Gal4/+>UAS-RelA/+, (9) repo-Gal4/+, (10) tub-Gal4/+>UAS-RelA/+, (11) tub-Gal4/+. Comparisons were based on two-way ANOVA and Bonferroni post-tests with **** denoting p < 0.0001. (E) Statistical analyses of SISL with RelA expressing in different subsets of PI. (1) wt, (2) M1KO, (3) UAS-RelA /+, (4) c767-Gal4/+ and UAS-RelA /+, (5) c767-Gal4/+ (6) Dh44-Gal4/+ and UAS-RelA /+, (7) Dh44-Gal4/+, (8) Dilp2-Gal4/+ and UAS-RelA/+,(9) Dilp2-Gal4/+, (10) SIFa2-Gal4/+ and UAS-RelA/+, (11) SIFa2-Gal4/+, (12) R19G10-Gal4/+ and UAS-RelA/+, (13) R19G10-Gal4/+. Comparison was based on t-test with *** denoting p < 0.001.

When *mesh1* gene was deleted in M1KO, nighttime SISL in M1KO flies was significantly more than that in wt flies, whereas daytime SISL were similar between the wt and M1KO mutant flies (Figure 8A and 8B). Although nighttime sleep latency during starvation night was still higher in M1KO (Figure S6A and S6B), there was no latency change after starvation during either night time or day time (Figure S6C and S6D). The M1KOGal4 knockin mutants were similar to M1KO flies in having exacerbated SISL (Figure 8C and S5). UAS-*mesh1* but not UAS-*mesh1E66A* could rescue the phenotype of SISL in mesh1 knockout flies, indicating that the ppGpp hydrolyzing activity is required for Mesh1 involvement in SISL. Expression of the bacterial ppGpp synthesizing enzyme RelA caused a decrease in SISL (Figure 8D), opposite to the SISL enhancing effect of Mesh1 (Figure 9A), indicating that ppGpp level, rather than any other unexpected activities of RelA or Mesh1 were involved.

**Figure 9.**
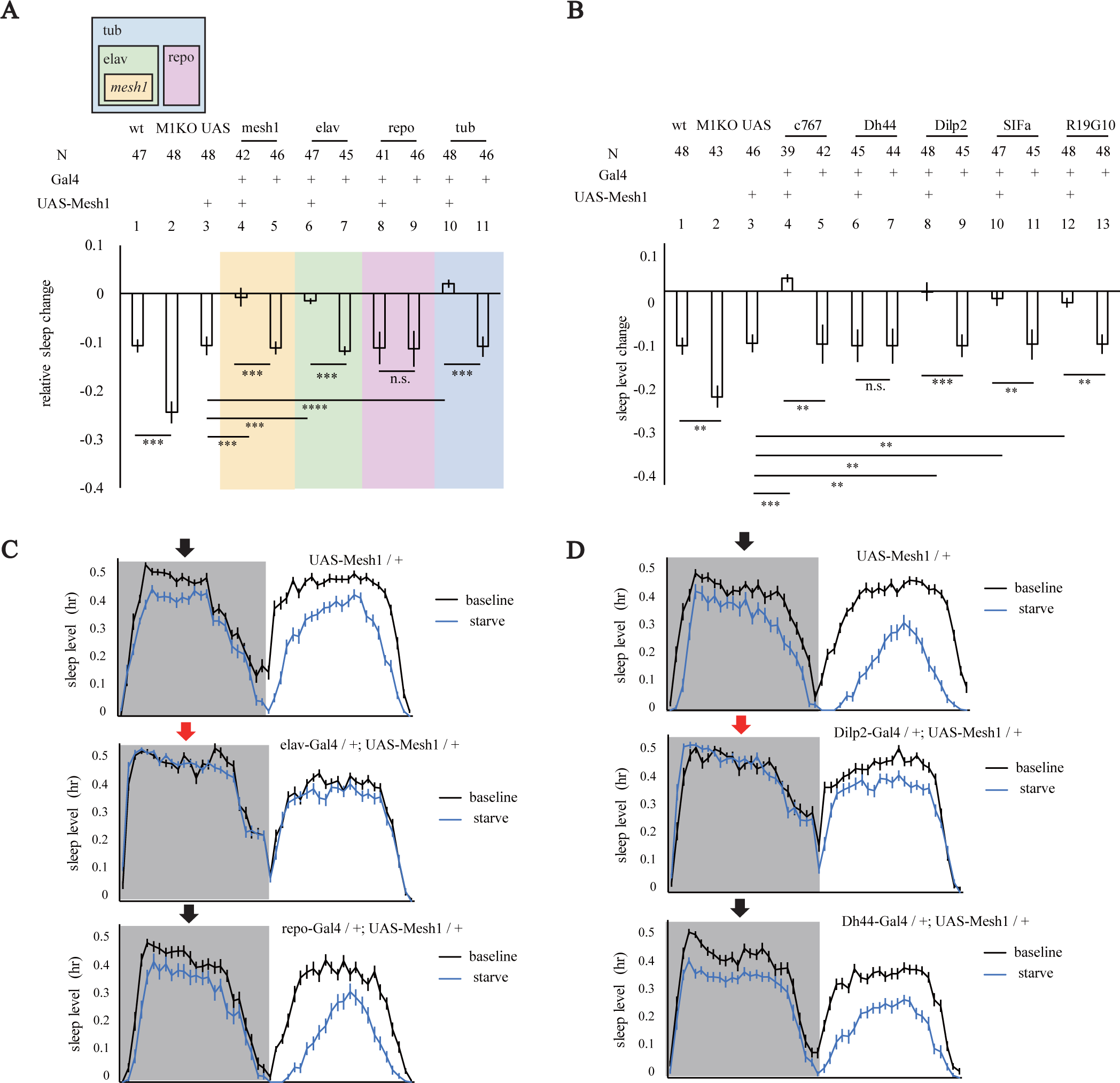
Effect of Mesh1 Overexpression of on SISL. (A) Statistical analyses of SISL with Mesh1 overexpression in neurons or other cells. (1) wt, (2) M1KO, (3) UAS-Mesh1/+, (4) M1KOGal4/+>UAS-Mesh1/+, (5) M1KOGal4/+, (6) elav-Gal4/+>UAS-Mesh1/+, (7) elav-Gal4/+, (8) repo-Gal4/+>UAS- Mesh1/+, (9) repo-Gal4/+, (10) tub-Gal4/+>UAS-Mesh1/+ tub-Gal4/+ (11). Comparisons were based on two-way ANOVA and Bonferroni post-tests with ***, **** denoting p < 0.001, p < 0.0001 respectively. (B) Statistical analyses of SISL with Mesh1 overexpression in different subsets of PI. (1) wt, (2) M1KO, (3) UAS-Mesh1/+, (4) c767-Gal4/+>UAS-Mesh1/+, (5) c767-Gal4/+, (6) Dh44-Gal4/+>UAS-Mesh1/+, (7) Dh44-Gal4/+, (8) Dilp2-Gal4/+> UAS-Mesh1/+, (9) Dilp2-Gal4/+, (10) SIFa2-Gal4/+>UAS-Mesh1/+, (11) SIFa2-Gal4/+, (12) R19G10-Gal4/+>UAS-Mesh1/+, (13) R19G10-Gal4/+. Comparison was based on t-test with *** denoting p < 0.001. (C) Representative sleep profiles of SISL in (A). Baseline sleep in black, and sleep during starvation in blue. Black arrows indicate that night SISL is not significantly different from wt; red arrows indicate SISL phenotype similar to that in M1KO. (D) Representative sleep profiles of SISL in (B).

SISL was increased by general or neuronal expression of RelA, but not by glial expression of RelA (Figure 8D, 9A and 9C). SISL was decreased by general or neuronal overexpression of Mesh1, but not by glial overexpression of Mesh1 (Figure 9A and 9C).

SISL was enhanced by RelA expression driven by Gal4 lines for c767, Dilp2, SIFamide or R19G10 (Figure 8E). SISL was reduced by Mesh1 overexpression driven by Gal4 lines for c767, Dilp2, SIFamide or R19G10 (Figure 9B, 9D). By contrast, Dh44 expressing neurons were not involved in ppGpp regulation of SISL because neither RelA expression nor Mesh1 overexpression in Dh44 neurons affected SISL (Figure 8E, 9B and 9D).

Thus, the same neurons required for ppGpp regulation of sleep latency and awakening are also involved in its regulation of SISL.

## DISCUSSION

We have discovered a new molecule in animals and provided evidence that it plays a physiological role. These results came from integrated genetic and chemical approaches. We have carried out a genetic screen of P element insertion lines which led to the discovery of a new mutation in the Drosophila ppGpp hydrolase Mesh1. Our chemical analyses indicate that ppGpp is present in Drosophila, not a contaminant of bacteria: germ free flies still had ppGpp and the content of ppGpp is regulated both by an enzyme encoded by a Drosophila gene and by starvation of flies. Our chemoconnectomic screen suggests involvement of PI neurons in ppGpp function. Further genetic intersection experiments confirm that ppGpp in specific PI neurons regulate sleep. Our findings indicate that ppGpp is not just a molecule present in prokaryotes and plants, but also exists and functions in animals. It regulates sleep, including sleep latency and SISL.

### Evidence for ppGpp Presence in Drosophila

We have shown that ppGpp is present in Drosophila and that it is hydrolyzed by Mesh1 which are expressed in specific neurons in flies. Moreover, ppGpp is detected in germ-free flies, indicating that ppGpp is synthesized in Drosophila (Figure 2G, 2H), not a contaminant of bacteria.

RSHs with the synthetase domain exist in both bacteria (Cochran and Byrne, 1974) and plants (van der Biezen *et al*., 2000), but not in animals (Atkinson *et al*., 2011; Sun *et al*., 2010). Our results suggest that the presence of a ppGpp synthetase in animals which does not contain an RSH like domain.

Plant RSHs are thought to result from lateral gene transfer events from bacteria (Ito *et al*., 2017; Field, 2018). Bioinformatic analyses and biochemical assays indicate that ppGpp synthetase homologs are distributed widely in plants including: dicotyledon A. thalaniana (van der Biezen *et al*., 2000), monocotyledon O. sativa (Xiong *et al*., 2001), green algae C. reinhardtii (Kasai *et al*., 2002), S. japonica (Yamada *et al*., 2004), N. Tabacum (Givens *et al*., 2004), pea plants (Takahashi *et al*., 2004), P. nil (Dabrowska *et al*., 2006a), C. annnum (Kim *et al*., 2009), and moss P. patens (Sato *et al*., 2015). Presence of the N-terminal chloroplast transit peptide (cTP), causes most plant RSHs to be located in the chloroplasts (Boniecka *et al*., 2017; Chen *et al*., 2014; Mizusawa *et al*., 2008; Takahashi *et al*., 2004; Sugliani *et al*., 2016; Sato *et al*., 2009; Takahashi *et al*., 2004). It is unclear whether a ppGpp synthetase exists in Drosophila mitochondria.

### Evidence for ppGpp Function in Drosophila

We obtain evidence for ppGpp function from four lines of experiments. First, three mutations in mesh1 caused sleep phenotypes in Drosophila. The first mutation is a P-element insertion (Eddison *et al*., 2012), whose phenotype (Figure 1A and 1B) and molecular nature we have characterized as *mesh1-ins* (Figure 1C). The second mutation is a knockout generated by us as M1KO (Figure 4A top panel). The third mutation is a knockin generated by us as M1KOGal4 (Figure 4A bottom panel). All three lines have the same phenotypes (Figure 1, 3 and 4).

Second, we have shown that the hydrolyzing activity of Mesh1 is required to rescue the *mesh1* knockout mutant phenotype: if a point mutation was introduced to amino acid residue 66 by converting it from E to A, then it was enzymatically inactive in vitro (Figure S1C and S1D) and unable to rescue the sleep phenotype in vivo (Figure 4B, 4C, 4D and 8C).

Third, expression of RelA, a bacterial ppGpp synthetase, in Drosophila phenocopied *mesh1* knockout mutants (Figure 6, 8D and 8E).

Fourth, the phenotypes of Mesh1 overexpression are opposite to those of RelA expression (Figure 7 and 9).

### Function of ppGpp in Neurons

Does ppGpp function in all cells or only in some cells? Results obtained here support that ppGpp functions in specific neurons in the PI to regulate sleep.

Mesh1 gene has been found to be expressed in specific neurons (Figure 5). Because Mesh1 is a ppGpp hydrolase, its expression can show the location where ppGpp is hydrolyzed, but not necessarily where it is synthesized or where it functions. When we used Gal4 lines from CCT to drive the expression of RelA, sleep phenotypes were observed in 6 lines which were all expressed in the PI (Figure 7). When Gal4 lines for PI neurons were used to drive RelA or Mesh1 expression, several of them could indeed cause sleep phenotypes (Figure 6, 7, 8 and 9). Expression of RelA or Mesh1 in glial cells did not affect sleep (Figure 6A, 6B, 7A and 7B).

Because it is difficult to imagine that both synthetase and hydrolase could cause the same multiple phenotypes in the same neurons if these neurons are not where ppGpp functions, our results are most consistent with the idea that ppGpp functions in these PI neurons.

### Molecular Targets of ppGpp

In bacteria, the best-known direct target of ppGpp is RNA polymerase (RNAP) (e.g., Artsimovitch *et al*., 2004; Barker *et al*., 2001; Kajitani and Ishihama, 1984; Kingston *et al*., 1981; Lindahl and Nomura, 1976). ppGpp interaction with the RNA polymerase leads to blockage of transcription initiation (Artsimovitch *et al*., 2004) and elongation (Kingston *et al*., 1981). There are also other targets for ppGpp (e.g., Paul *et al*., 2005); Gourse *et al*., 2018; Maciag *et al*., 2010; Nomura *et al*., 2014; Sherlock *et al*., 2018; Corrigan *et al*., 2016; Pao and Dyess, 1981; Dalebroux and Swanson, 2012; Zhang *et al*., 2019). A recent study utilizing a ppGpp-coupled bait uncovered new ppGpp target proteins in bacteria, including a large group of GTPase and metabolism-related enzymes (Wang *et al*., 2019). Future studies are required to identify the molecule(s) directly mediating the sleep regulating function of ppGpp in Drosophila.

## SUPPLEMENTAL INFORMATION

Supplemental information includes 6 Figure.

## ACKNOWLEDGEMENTS

We are grateful to Dr. U. Heberlein for sharing P-element insertion mutant library, Drs. J. Ni and G. Gao for providing us with CRISPR/Cas9 plasmids and flies, to Dr. Z.-F. Gong for sharing antibodies, to the Beijing Commission of Science and Technology and the National Natural Science Foundation of China (Project 31421003 to Y. R.) for grant support.

## AUTHOR CONTRIBUTIONS

Y.R supervised and initiated the project. W. Y. performed the majority of experiments and data analysis. X. H. W. carried our experiments of UPLC-MS. W. Y., E. Z. and H. D. X. carried out experiments of sleep screen on P-element insertion mutants. W. Y. implemented the fly tracing program, and E. Z. developed software for sleep analyses. W. Y. and Y. R. wrote the manuscript.

## DECLARATION OF INTERESTS

The authors declare no competing interests.

---

**Figure S1.**
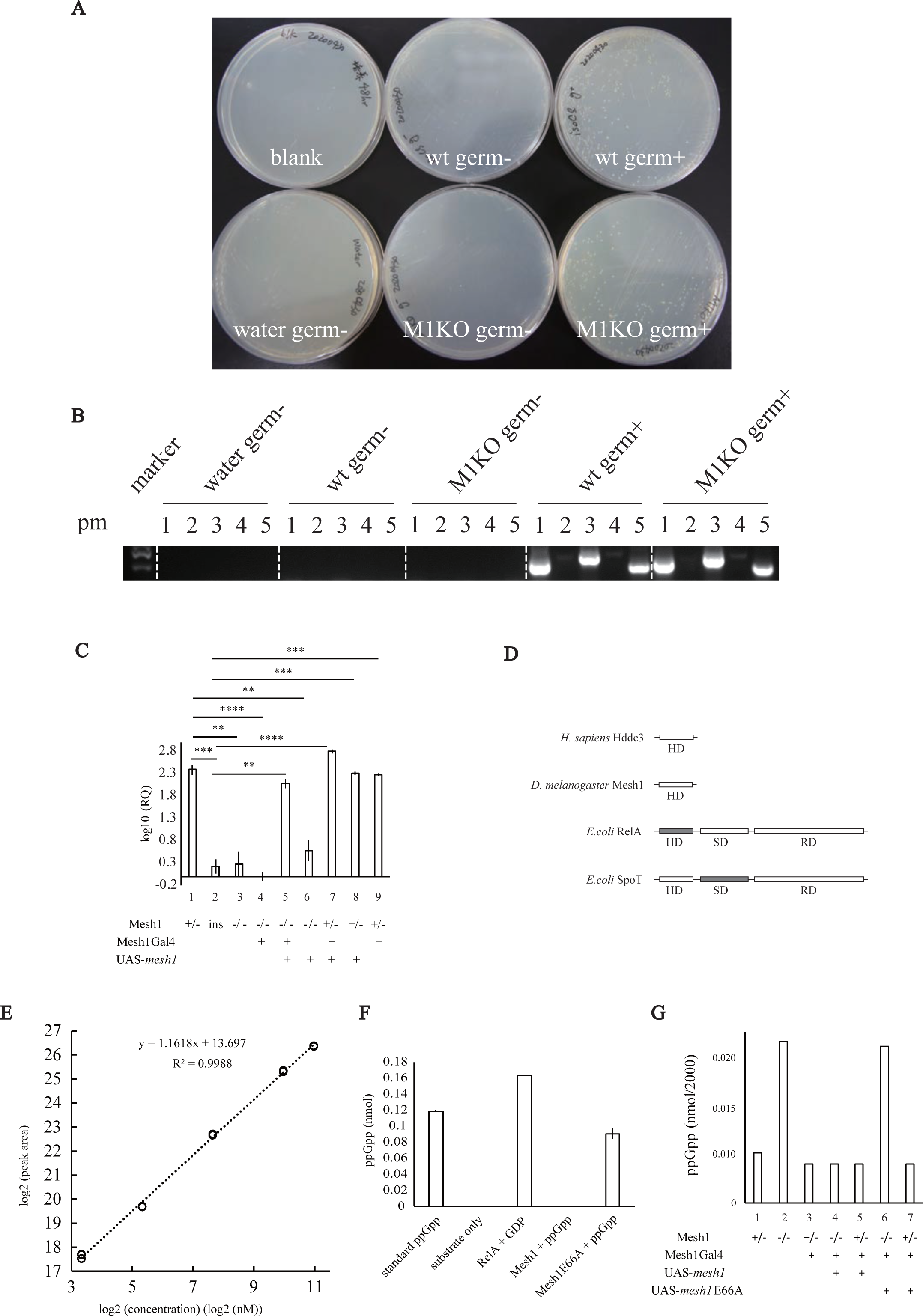
Quantifications: Mesh1 RNA and ppGpp. (A) Examination of germ-free status by growing bacteria on LB plate. 6 conditions were examined: (1) blank: direct incubation of LB plate; (2) water germ-: strike LB plate with germ-free water; (3) wt germ-: extract germ-free wt flies with germ-free water, and strike LB plate with the extract; (4) M1KO germ-: extract germ-free M1KO flies with germ-free water, and strike LB plate with the extract; (5) wt germ+: extract normal wt flies with germ-free water, and strike LB plate with the extract; (6) M1KO germ+: extract normal M1KO flies with germ-free water, and strike LB plate with the extract. Among these conditions, bacteria only appear on plates of wt germ+ and M1KO germ+. (B) Examination of germ-free status by 16S rDNA PCR. 5 pairs of 16S PCR primers (denoted as pm) were used, including (1) 16S-27F/16S-519R; (2) 16S-357F/16S-907R; (3) 16S-530F/16S-1110R; (4) 16S-926F/16S-1492R; (5) 16S-1114F/16S-1525R. The upper and lower bands at marker lane (Trans2K) correspond to 500bp and 250bp. PCR signals by pm1, pm3 and pm5 were detected in both controls (wt germ+ and M1KO germ+), while no signals were detected in all three germ-free groups (water germ-, wt germ-, M1KO germ-). (C) Measurement of Mesh1 RNA by quantitative polymerase chain reaction (qPCR). Log_10_(RQ) is the log form of relative quantity of Mesh1 mRNA in Drosophila. (1) M1KO/+, (2) *mesh1*-ins, (3) M1KO, (4) M1KOGal4/M1KO, (5) M1KOGal4/M1KO and UAS-Mesh1/+, (6) M1KO/M1KO and UAS-Mesh1/+, (7) M1KOGal4/+, UAS-Mesh1/+, (8) M1KOGal4/+ and UAS-Mesh1/UAS-Mesh1, (9) M1KOGal4/+. (D) Diagrams of the ppGpp synthetase domain (SD) and the hydrolase domain (HD) in RSH proteins from E. coli, D. melanogaster, and H. sapiens. The mammalian hddc3 and the Drosophila Mesh1 proteins contain only HD, the bacterial RelA protein contains a weak HD, an active SD and a regulatory domain (RD). Bacterial SpoT contains an active HD, a weak SD and a RD. (E) Standard curve of MS peak area to ppGpp concentration. x-axis is log2 value of standard ppGpp concentration (nM), and y-axis is log2 value of peak area of MS signal. (F) Measurement of ppGpp in vitro. Blank is the mixture of reaction buffer, GDP and ATP. Standard ppGpp is a commercial sample. RelA+GDP is the addition of purified RelA to the mixture of reaction buffer, GDP and ATP. Mesh1+ppGpp is the standard ppGpp plus purified Mesh1. Mesh1E66A+ppGpp is the standard ppGpp plus the mutant protein Mesh1E66A expressed in *E. coli* and purified. (G) Measurement of ppGpp level in vivo. (1) M1KO/+, (2) M1KO, (3) M1KOGal4/+, (4) M1KOGal4/M1KO and UAS-Mesh1/+, (5) M1KOGal4/+ and UAS-Mesh1/+, (6) M1KOGal4/M1KO and UAS-Mesh1E66A/+, (7) M1KOGal4/+ and UAS-Mesh1E66A/+.

**Figure S2.**
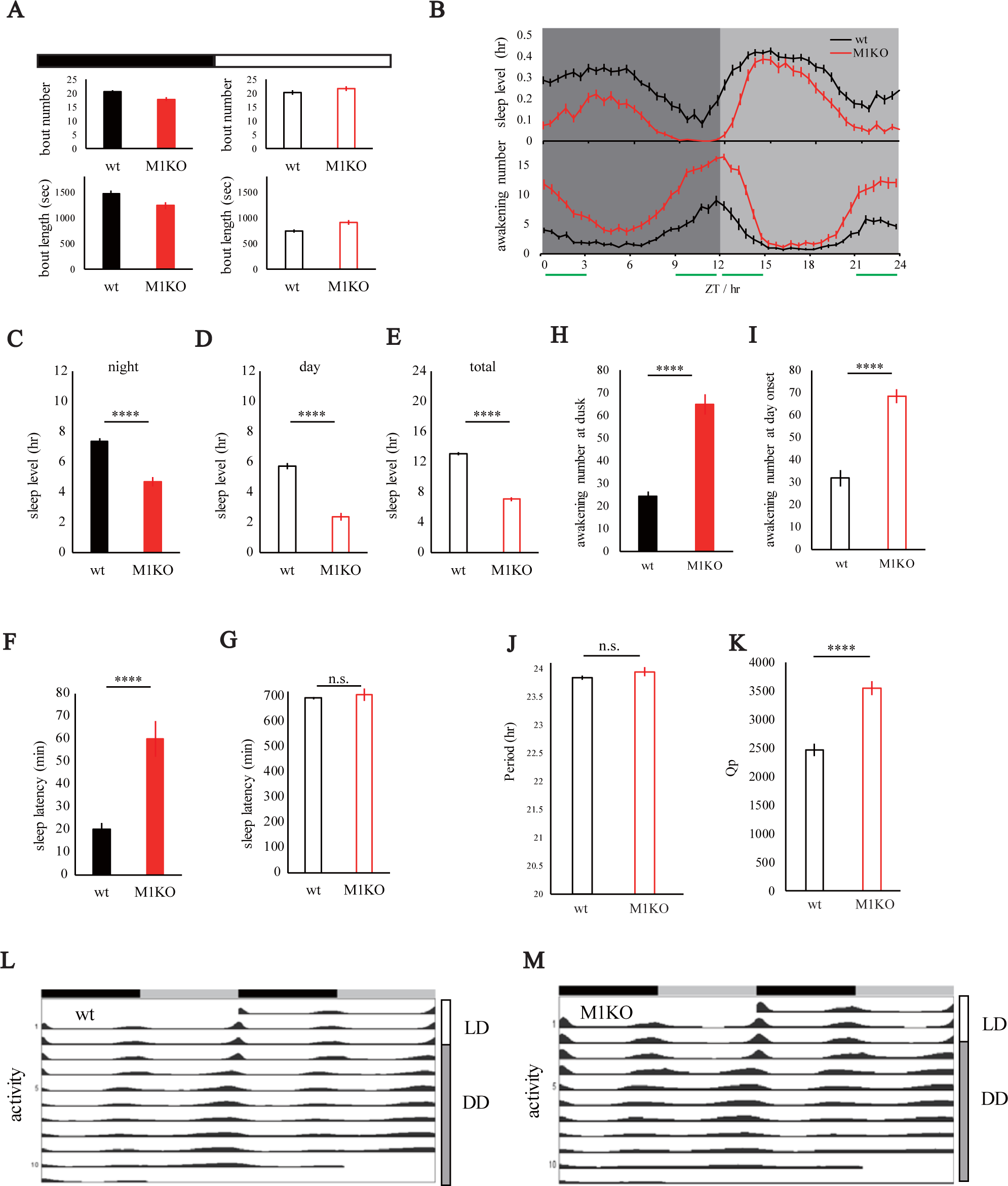
Phenotypic Analyses of Sleep and Circadian Rhythm. (A) Sleep analysis under LD condition. Nighttime bout number and bout length (in filled boxes on the left) and daytime bout number and bout length (unfilled boxes on the right) were not significantly different between the wt and the M1KO flies. (B) Sleep and awakening number profiles under DD condition. (C-I) Sleep analyses under DD conditions. Sleep level at presumptive night (C), at presumptive day (D), total sleep level (E), awakening number at dusk (H), at dawn (I), sleep latency at presumptive night (F), at presumptive day (G). (K-L) Periodic length (K) and rhythmicity strength Qp (L) of wt and M1KO flies. (L) Circadian profiles of wt flies under the LD and DD conditions. (M) Circadian profiles of M1KO flies under the LD and DD conditions.

**Figure S3.**
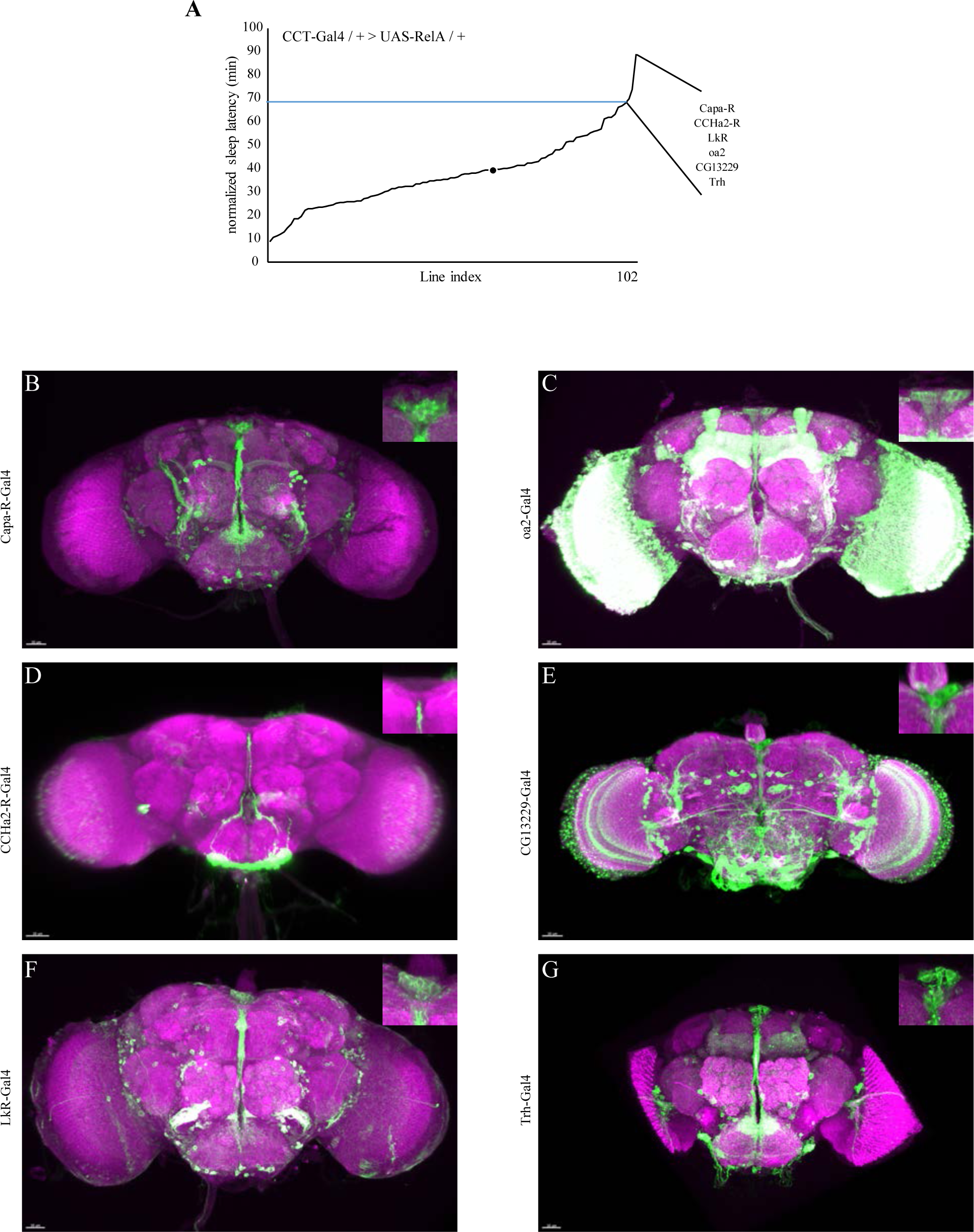
A Functional Screen of CCT-Gal4 Lines. (A) A pilot screen of sleep latency with 102 CCT-Gal4 lines driving RelA expression. Candidates above blue line showing three standard deviations away from the mean. 6 Gal lines were found to be able to increased sleep latency significantly: Capa-R, CCHa2-R, LkR, OA2, CG13229 and Trh. (B-G) mCD8-GFP expression driven by each of the CCT Gal4 lines: Capa-R (B), OA2 (C), CCHa2-R (D), CG13229 (E), LkR (F) and Trh (G). PI neurons were observed in all these lines.

**Figure S4.**
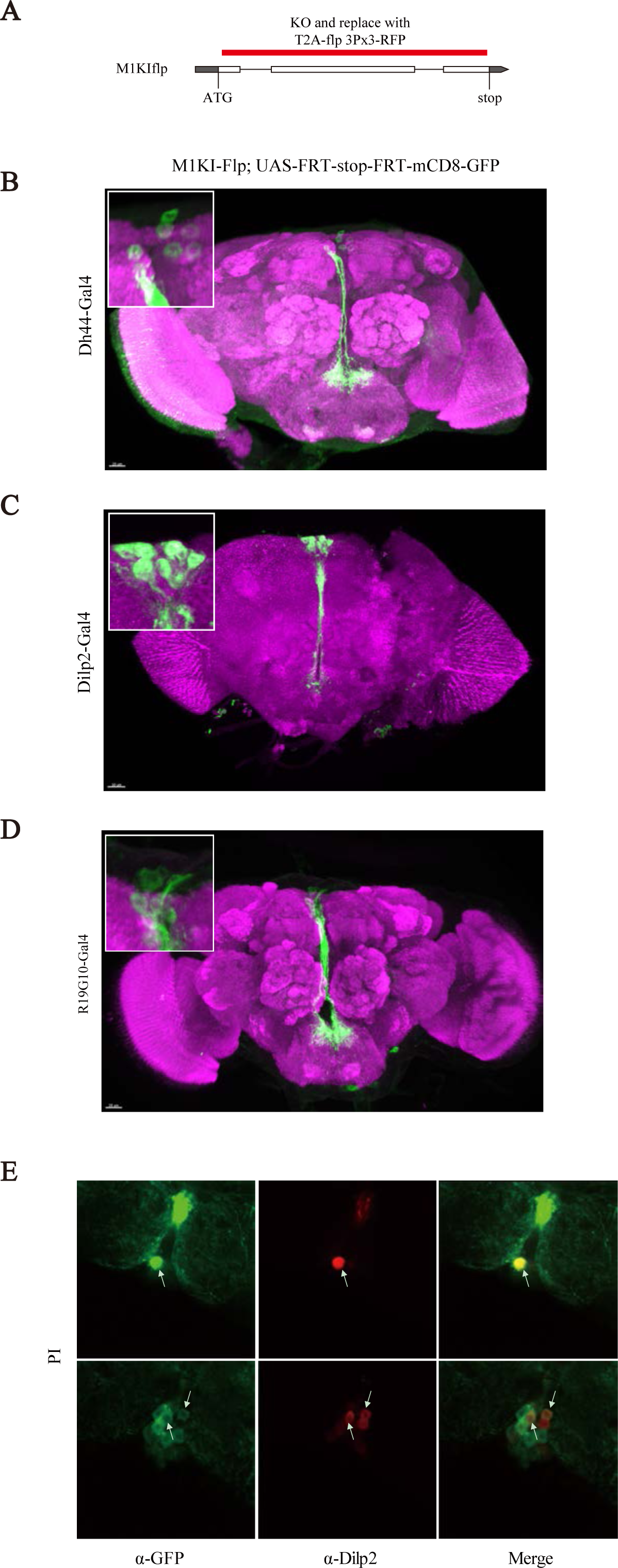
Expression Examined by Intersection with Mesh1. (A) Diagram of M1KIflp. The CDS after the start codon of *mesh1* gene was replaced by 2A-flp-3Px3-RFP. This is similar to Mesh1Gal4 but with Flp in place of Gal4. This line was used to intersect with Gal4 lines from the CCT screen (Capa-R, CCHa2-R, LkR, OA2, CG13229 and Trh). Each was indeed expressed in Mesh1 positive PI neurons. Shown from B to D are examples. (B-D) The UAS-FRT-stop-FRT-mCD8-GFP expression patterns of M1KIflp and PI-Gal4 intersection lines. Insets show higher magnification views of the PI. (E) An anti-Dilp2 antibody was used in immunostaining in M1KOGal4>UAS-mCD8-GFP flies and showed Mesh1 positive neurons to be also positive for Dilp2.

**Figure S5.**
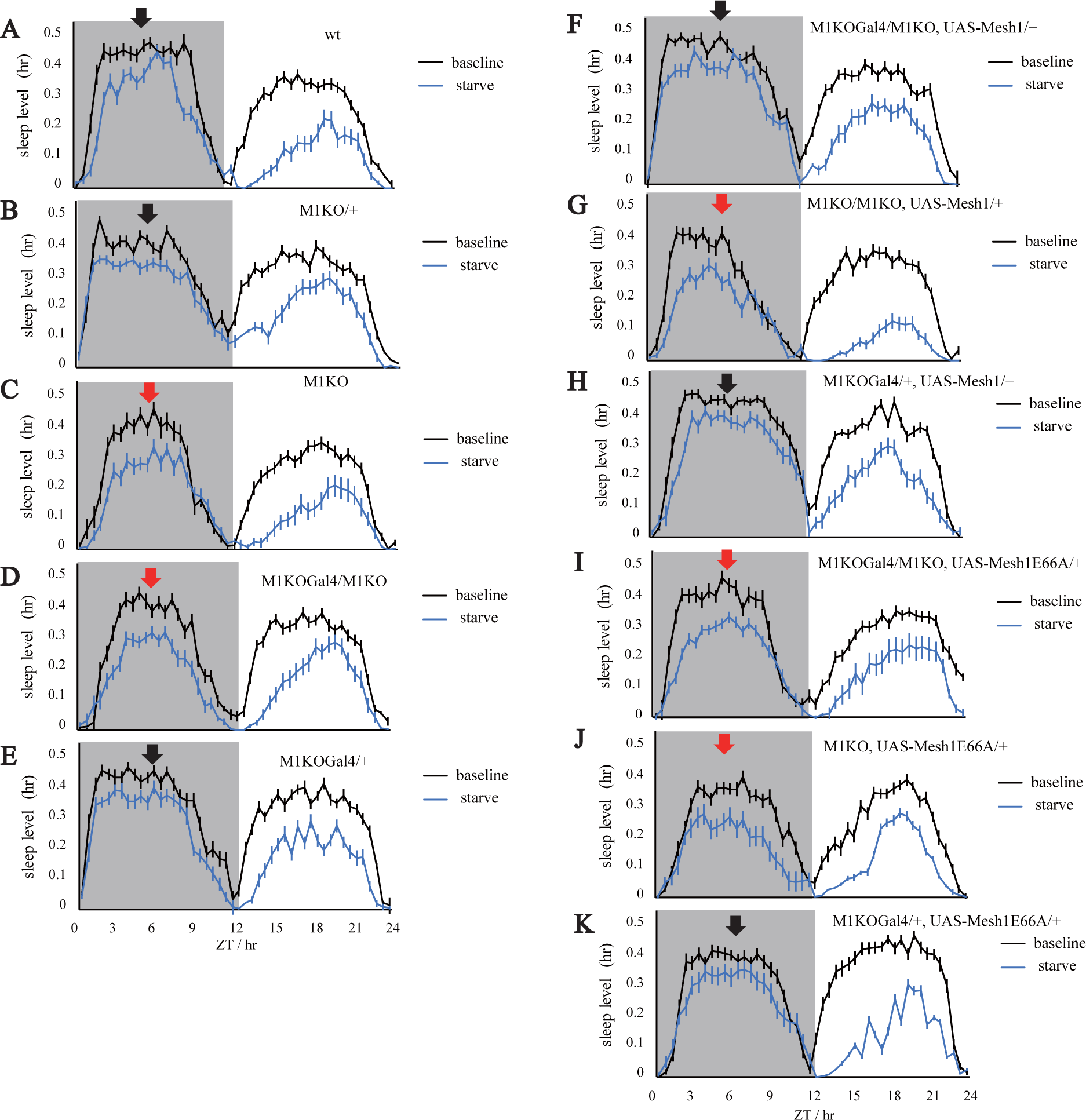
Sleep profiles of rescue lines in SISL (A-K) baseline sleep is shown in black, and sleep during starvation shown in blue; black arrows indicate that night SISL is not significantly changed compared to wt or corresponding parental controls; red arrows indicate that a specific genotype phenocopied M1KO in SISL.

**Figure S6.**
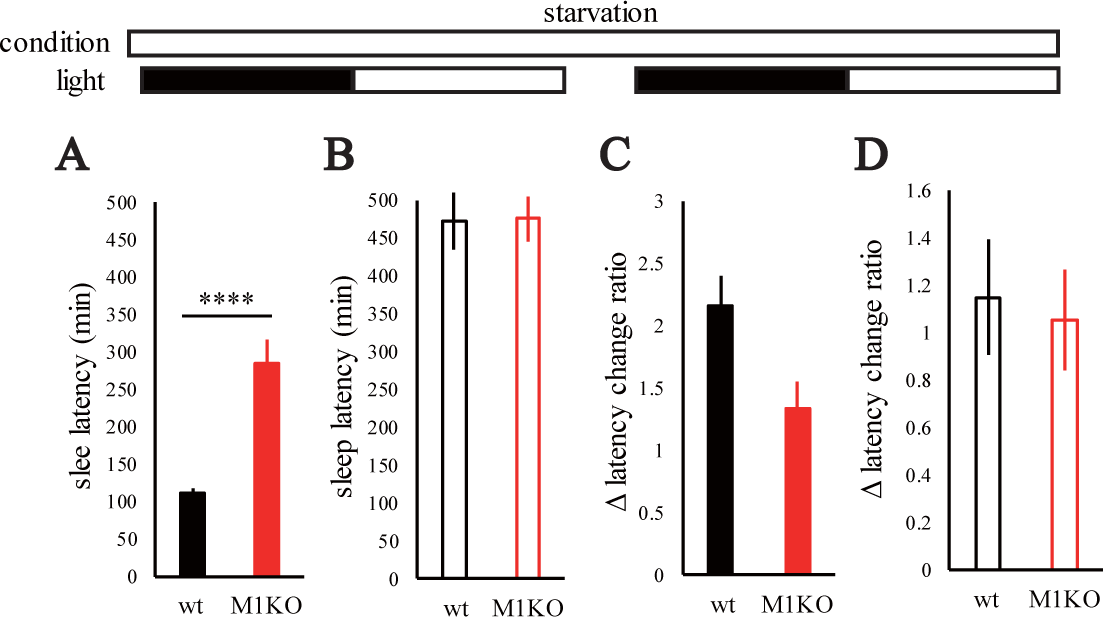
Sleep Latency in SISL. Sleep latency (A, C) at night or day (B, D). A and B are before starvation, whereas C and D are during starvation.

Supplementary Table S1. Primers for PCR, qPCR and Genotyping

Supplementary Table S2. Standard Curve Data of MS peak area to ppGpp concentration.

## EXPERIMENTAL PROCEDURES

### Fly Stocks and Rearing Conditions

All flies were reared on standard corn meal at 25℃ and 60% humidity, under 12hr:12hr LD cycle (unless specified otherwise). Before behavioral assays, stocks were backcrossed into the background of an isogenized Canton S wt line in the lab for 7 generations (Zhou, Rao and Rao, 2008).

Lines ordered from the Bloomington Stock Center included: 47887 (R19G10-Gal4), 30848 (c767-Gal4), 51987 (Dh44-Gal4), nos-phiC31, 37516 (Dilp2-Gal4), 5137 (UAS-mCD8-GFP). SIFa-Gal4 was a gift from Dr. J. Veenstra (University of Bordeaux). P-element insertion lines were a gift from Dr. U Heberlein, and CCT-Gal4 library was a collection previously generated in our lab (Deng *et al*., 2019). isoCS and w^1118^ were wild-type and white-eye wild-type lines.

### Reagents and Plasmids

PCR was performed with Phanta-Max Super-Fidelity DNA Polymerase (Vazyme). Genotyping PCR was performed with 2x Taq PCR StarMix with Loading Dye (GenStar). Restriction enzymes KpnI-HF, SacII, NotI-HF, XbaI, XhoI, BamHI-HF, DpnI, EcoRI-HF and XbaI were from New England BioLabs. Total RNA was extracted from flies with RNAprep pure Tissue Kit (TIANGEN). Reverse transcription for cDNA cloning was performed with PrimeScript^TM^ II 1^st^ Strand cDNA Synthesis kit (Takara), Gibson assembly was performed with NEBuilder HiFi DNA

Assembly Master Mix. Transformation for cloning was performed with Trans-5α (TransGen), and transformation for expression was performed with Transetta (TransGen). BL21 (TransGen) was used as the bacterial gene template. Reverse transcription for quantitative PCR (qPCR) analyses was performed with PrimeScript RT Master Mix kit (Takara). qPCR was performed with TransStart Top Green qPCR SuperMix kit (TransGen).

pACU2 was a gift from Drs. Lily and YN Jan. The plasmids previously used in our lab included: pBSK, pET28a+. Templates of STOP-attP-3Px3-RFP, T2A-Gal4-3Px3-RFP, and T2A-flp-3Px3-RFP were previously generated in the lab (Deng *et al*., 2019).

### Molecular Cloning and Generation of Genetically Modified Flies

Generation of all KO and KI lines was based on the CRISPR-Cas9 system with homologous recombination, according to previous procedures described (Ren *et al*., 2013). U6b vector was used for the transcription of sgRNA (Ren *et al*., 2013) and construction of targeting vectors were based on previous procedures in our lab (Deng *et al*., 2019).

To generate KO lines, a mixture of two U6b-sgRNA plasmids and one targeting vector was injected into Drosophila embryos. To generate U6b-sgRNA, two sgRNAs were selected on website (https://www.flyrnai.org/crispr), and designed into a pair of primers (M1KOSgRNA-F and M1KOSgRNA-R) without PAM sequence. The other pair of primers (U6b-laczrv and U6b-primer1) were designed on the backbone of U6b vector, so that they could generate PCR products containing two parts of U6b (shorter fragment by M1KO sgRNA-F and U6b-laczrv, longer fragment by M1KO sgRNA-R and U6b-primer1), which shared overlapping sequences in both ends. U6b-sgRNA plasmid was built by Gibson-assembly of the PCR products. To generate the targeting vector, two fragments (2kbps) flanking the entire CDS of gene *mesh1* except start codon were cloned as 5’Arm (PCR with primers M1KO5F and M1KO5R) and 3’Arm (PCR with primers M1KO3F and M1KO3R). Vector pBSK was digested with *KpnI* and *SacII*, and the PCR products were introduced into the digested pBSK by Gibson-assembly. New restriction sites (*NotI* and *XhoI*) were introduced between the two arms for further use. To generate a targeting vector for M1KO, the fragment STOP-attP-3Px3-RFP was cloned with primers attP2M1KOF and attP2M1KOR, and inserted between *NotI* and *XhoI* by Gibson-assembly; and for M1KOGal4, T2A-Gal4-3Px3-RFP was inserted at same location. A mixture of U6b-sgRNA and the targeting vector was injected into embryos of *nano*-Cas9 or *vasa*-Cas9 (Ren *et al*., 2013). F1 individuals of RFP+ eyes were collected after being crossed with w^1118^ flies.

To generate KI line M1KIflp, similar procedures were performed. For U6b-sgRNA, primers for shorter fragment were M1KISgRNA-F and U6b-laczrv, while primers for longer fragment were M1KIsgRNA-R and U6b-primer1. For the targeting vector, PCR for 5’Arm was performed with M1KI5F and M1KI5R, and 3’Arm with M1KI3F and M1KI3R. The site flanked by two arms was selected near the very end of *mesh1* CDS, so that the stop codon was removed and the CDS was fused with T2A-flp-3Px3-RFP with primers flp2M1KIF and flp2M1KIR.

Generation of transgenic UAS lines was based on vector pACU2 (Han, Jan and Jan, 2011). Drosophila cDNA was generated by reverse transcription from total RNA. By using the cDNA as template, *mesh1* CDS was amplified with primers (M1CDSF and M1CDSR), and inserted into digested pBSK (by *EcoRI* and *KpnI*). Point mutation of *mesh1*E66A was generated on this plasmid with primers M1E66AF and M1E66AR. Both *mesh1* CDS and Mesh1E66A were cloned with primers M1ACU2F and M1ACU2R. RelA sequence was cloned from BL21 bacteria with primers RAACU2F and RAACU2R. All the above were inserted into digested pACU2 (*EcoRI* and *XbaI*) by Gibson assembly. pACU2 constructs were inserted into attP2 by nos-phiC31 during embryo injection.

All fly stocks were confirmed by PCR and qPCR, and primers were shown in Supplementary Table S1.

### Molecular Cloning and Inducible Expression in Bacteria

pET28a+ was used for bacterial expression of Mesh1 and RelA proteins. *mesh1* CDS and *mesh1*E66A were generated with primers M1ET28F and M1ET28R. RelA was cloned from BL21 bacteria with primers RAET28F and RAET28R. All the above were inserted into pET28a+ by Gibson assembly. Competent cells transformed with the above constructs were amplified for inducible expression. To induce expression, a colony was inoculated into 1ml kanamycin containing Luria broth (LB) media, and incubated at 37℃ for 2 hr. 800μl was transferred into a flask with 800ml kanamycin+ LB media and incubated for ∼3hr at 37℃ 220rpm, until OD_600_ was 0.5 to 0.6. 800μl 1M isopropyl β-D-1 thiogalactopyranoside (IPTG) was added into the flask. For RelA, the induction was at 37℃ for 3hr; for Mesh1 and Mesh1E66A, it was at 16℃ for 16 hr. Bacteria was harvested by centrifugation followed by lysis with ultrasonication (Power: 30%, lysis 2s, wait 2s, 30 cycles) and centrifugation. Supernatants were passed through nickle columns, followed by two times of wash with 10ml binding buffer (20mM Tris-HCl pH 7.4, 0.5M NaCl, 5mM imidazole), and elution with 5ml elution buffer (20mM Tris-HCl pH7.4, 0.5M NaCl, 500mM imidazole). Each step was monitored on 10% SDS-PAGE to check protein expression. Eluted protein was enriched in Millipore Amicon Ultra-15, and re-suspended in 1ml of protein storage buffer (20mM Tris-HCl pH7.4, 150mM NaCl, 0.3% CHAPS, 1mM DTT).

### Generation of Germ-Free Flies

Replace the sponge of bottle containing standard fly medium with autoclavable PP bag, and autoclave twice. Evaporate water of food surface in laminar hood and under UV light for 1 hour.

According to previous report (Kietz, Pollari and Meinander 2018), prepare a simple device for Drosophila egg collection and filter for egg wash, and use the device to collect fly eggs for 5 hours, while filters were soaked in 75% EtOH overnight.

Prepare 5x 50ml centrifuge tubes, filled with the following solutions: A. 3.3% Walsh hand-sanitizer; B. 75% EtOH; C. 1% active chlorine; D. 0.1% TritonX-100 + 1xPBS; 1. E. sterilized ddH2O.

Use cell scraper to collect eggs onto filters, and wash filters from A to E, and disperse the resuspended eggs onto autoclaved fly medium.

Keep rearing at clean incubator for 20 days, and amplify the generated germ-free flies.

### Extraction and Measurement of ppGpp

To extract ppGpp from *Drosophila*, 24ml formic acid was added into about every 2000 flies. After grinding 10,000 rpm for 15 seconds, 3ml 30% tri-chloric acetic acid was added to precipitate proteins. The above procedures were repeated to collect extracts if necessary. Supernatants were enriched with SPE column (Waters Oasis 6cc WAX), and effluent was titrated and lyophilized overnight, and powders were re-suspended with 200μl water.

To measure the level of ppGpp, ZORBAX Eclipse XDB-C18 2.1x100mm column was used in UPLC-MS (Thermo ULtiMate 3000, and Thermo QE HF-X), according to a modified version of previous method (Ihara, Ohta and Masuda, 2015). The mobile phase included: Buffer A (8mM *N,N*-dimethylhexylamine, 160μl acetic acid, 500ml water) and Buffer B (Acetonitrile). The UPLC program was: at 0min, A:B = 100%:0%; at 10min, A:B = 40%:60%, with linear increment; at 10.01-14min, keep A:B = 0%:100%; at 14.01-18min, keep A:B=100%:0%. The m/z of ppGpp should be 601.95, and fly number was used for normalization.

### Behavioral Assays

To analyze baseline sleep under 12hr:12hr LD cycles, approximately 48 flies of each genotype were loaded into glass tubes for video tracing (fps=1), which was analyzed by an in-house software as described previously (Qian *et al*., 2017; Dai *et al*., 2019; Deng *et al*., 2019). Continuous immobility of >5min was defined as a sleep bout (Hendricks *et al*., 2000, Shaw *et al*., 2000). Sleep latency at dusk was defined as the that from light-off to the point when the 1^st^ sleep bout appeared.

To analyze the circadian rhythm, activities under 12hr:12hr LD cycles were recorded for 7 days, which was then computed by ActogramJ (Schmid *et al*., 2011). Period was calculated by periodogram of Chi-square, and rhythmicity was assessed with Qp.

To analyze awakening numbers, video tracing data was converted to data of simulated cross-beam. In the simulation, the middle line for each tube was set as the virtual beam. According to a previous study (Tabuchi *et al*., 2018), brief awakening was defined as 1 cross per min, and the awakening number was the sum of such events in every 30 min. Awakening number of dusk was the sum of brief awakenings at ZT0-3, and awakening number of dawn was the sum of brief awakenings at ZT9-12.

To test starvation-induced sleep loss (SISL), sleep was recorded for the first 3 days (3rd day defined as baseline), and flies were quickly transferred to 1% agar at the end of the light phase of 3^rd^ day, followed by a 24 hr recording of sleep during starvation. SISL ratio was defined as (starvation sleep-baseline sleep)/ baseline sleep.

### Immunohistochemistry and Imaging

To prepare flies for imaging, 5 flies per genotype were dissected in phosphate-buffered saline (PBS). Dissected tissues were transferred to a tube of 400μl 2% PFA, and fixed for 55 min. Tissues were washed 3 times with 400μl brain wash buffer (PBS containing 1% TritonX-100, 3% m/V NaCl), and then transferred to 400μl blocking buffer (PBS containing 2% TritonX-100, 10% normal goat serum), followed by incubation at 4℃ overnight. Tissues were transferred to dilution buffer (0.25% TritonX-100, 1% NGS, 1x PBS) and added with primary antibodies including, 1:1000 chick anti-GFP (Abcam) and 1:40 mouse anti-nc82 (DSHB). Tissues were stained at 4℃ overnight, followed by 3 washes with 400μl brain wash buffer. Samples were transferred to fresh dilution buffer containing secondary antibodies including, 1:200 Alexa Fluor goat anti-chick 488 (Invitrogen) and 1:200 Alexa Fluor goat anti-mouse 633 (Invitrogen), followed by 3 washes with 400μl brain wash buffer. To prepare for imaging, samples were placed into a drop of Focus Clear (Cell Explorer Labs, FC-101), which was restricted by a piece of paper circle on a glass slide. Samples were visualized on Zeiss LSM 710 confocal microscope.

## CONTACT FOR REAGENT AND RESOURCE SHARING

Further information and requests for resources and reagents should be directed to and will be fulfilled by the Lead Contact, Yi Rao.

## Notes

### Competing Interest Statement

The authors have declared no competing interest.

